# In-slide sample preparation allows for rapid triple staining and the detection of specific cell wall components in monocot roots

**DOI:** 10.64898/2026.02.13.705847

**Authors:** Julia Zheku, Raju Soolanayakanahally, M. Arif Ashraf

## Abstract

Anatomical and histochemical imaging of monocot root systems relies on tissue sectioning and cell wall staining dyes because molecular reporter lines are limited for most organisms. Distinct staining dyes require variable incubation time and concentration across different tissues and organisms. As a result, staining with multiple dyes becomes time consuming or challenging. Here, we report a rapid method to perform simultaneous triple staining and clearing of samples on a glass slide. The entire protocol requires ∼4 hours and a smaller volume of stain than traditional methods. We tested this method using the roots of 3 crops, *Triticum aestivum* (wheat), *Zea mays* (maize), and *Cucurbita pepo* (winter squash), and the stems of 4 crops, *Cucurbita pepo* (winter squash), *Solanum lycopersicum* (tomato), *Tagetes patula* (marigold), and *Camelina sativa* (false flax). We have demonstrated the presence of exodermis in wheat roots and a multiseriate exodermis in maize roots. Images produced with this method are compatible with ImageJ/Fiji based quantification methods. This sample preparation method ultimately simplifies the staining process while empowering a quantitative cell biology approach to study cell-type specific distribution of cell wall materials in grass root systems and beyond.

## 1. Background

In the field of plant cell biology, finding clear, robust material to examine and quantify is usually the first problem that needs solving. Plant cells can at times be difficult to study under the microscope due to autofluorescent pigments and presence of polysaccharides like cellulose in plant cell walls (Piccinini et al., 2024). When working with roots of the model organism *Arabidopsis*, visualization is straightforward and provides clear images, due to relatively thin tissue (Truernit et al., 2008) However, many discoveries at the cellular level do not align between *Arabidopsis* and other economically important crops because of major anatomical differences (Roeder et al., 2025). Certain re-enforced root tissues, like the exodermis (Cantó-Pastor et al., 2024; Chen et al., 2022; Kreszies et al., 2018) and multiple cortex layers (Li et al., 2026; Ortiz-Ramírez et al., 2021; Roeder et al., 2025), are simply not present in *Arabidopsis*, which limits our understanding. Root system architecture and individual root morphology are also significantly variable between eudicots, such as *Arabidopsis*, and monocots, such as wheat and maize (Viana et al., 2022). Comparatively, less is known about monocot crop roots as the plants themselves are noticeably larger and harder to visualize at the cellular level. Such disparities highlight the importance of researching *Arabidopsis* and crop plants, especially monocots, in tandem.

In order to study monocot root anatomical features, sectioning and staining of thick tissues is required due to the lack of readily available fluorescent protein marker lines. Within such investigations, classification of tissues and determination of specific cell wall compounds are major goals. Combinations of vibratome, microtome, ultramicrotome and hand sectioning are generally used, yielding sections that are on average 50-250 µm thick (Verhertbruggen et al., 2017). Both bright field and fluorescence microscopy have been used to define distinct cell wall compounds and tissues in monocot roots. For instance, suberin and lignin can be visualized by Sudan III and phloroglucinol/HCl staining, respectively (Jensen, 1962; Ouyang et al., 2020; Spicher, 1985). Sudan III and phloroglucinol/HCl staining result in orange and bright red/pink stains, which can be observed using a bright field microscope. In contrast, lignin can be seen with berberine hemisulphate staining which shows yellow fluorescence under UV light (Tylová et al., 2017). Finally, fluorescent dyes, such as, but not limited to Calcofluor White, Auramine O, SCRI Renaissance 2200, Basic Fuchsin, Fluorol Yellow 088, Aniline Blue, and Direct Red 23 display specific binding capacity with cell wall materials like cellulose, suberin, lignin, callose, and cutin (Kitin et al., 2020; Piccinini et al., 2024; Ursache et al., 2018). Given the resources available, an excitation of either UV (around 275–405 nm) or blue (around 450–490 nm) light coupled with an emission filter capable of detecting 400–550 nm (blue green) works very efficiently for visualizing key tissues and monocot root cell wall components using epifluorescence and confocal microscopes.

Quantification of root features and microscopic images can be uniquely finicky and unreliable. Many methods utilize either the binary method (presence or absence of staining) or a scoring system (a range of values arbitrarily assigned based on the fluorescence or staining intensity) (Enstone and Peterson, 2005; Ouyang et al., 2020). Such quantification is prone to human error and bias. Moreover, autofluorescence is widely used and convenient for observing root morphology and counting cortical cell layers (Baca Cabrera et al., 2026; Ortiz-Ramírez et al., 2021; Zhang et al., 2025). Relying on autofluorescence comes with limitations because high background or spectral overlap from a range of plant fluorophores eliminates the ability to directly identify polymers, making it unreliable for quantitative imaging approaches (Czymmek et al., 2025; Donaldson, 2020). Quantitative imaging, based on histochemical staining, offers an unbiased approach, as it relies on pixel data of the experimental image to calculate a numerical value for fluorescence intensity. The combination of multiple staining dyes, which are component specific and visualized at different fluorescence channels (Piccinini et al., 2024; Sexauer et al., 2021) allows for the quantification and comparison of specific cell wall components within a specimen or across many samples, immensely increasing quantitative power.

Current sample preparation methods provide adequately stained samples for visualization and quantification yet are labor intensive and time consuming. Firstly, many protocols require a clearing step, which utilizes organic solvents to clear endogenous autofluorescent compounds that interfere with imaging (Kurihara et al., 2015). This distinct step is dependent on the given sample type and can take up to one week, in the case of extremely woody tissues (Supplemental Table 1). Furthermore, staining and washing steps are typically conducted in well plate dishes or falcon tubes and require continual movement of samples. This process is inconvenient when working with small samples and also promotes mechanical damage to delicate root tissues. Additionally, stains such as Sudan III and phloroglucinol/HCl, are limited to one staining solution at a time, yielding a single output color. While this kind of staining is more accessible at times, it limits the number of cell wall components visualized in the same sample. These problems can be resolved by 1) dissolving stain powders into a clearing solution, in practice effectively clearing and staining tissue at the same time, 2) minimizing the movement of delicate samples, and 3) using multiple staining dyes to increase quantitative power. Several protocols have already demonstrated the power of simultaneous staining and clearing, in other words dissolving staining dyes directly into a clearing solution (Manzano et al., 2025; Sexauer et al., 2021)(Supplemental Table 1).

To further streamline the staining process, we are reporting the in-slide sample preparation method. In this protocol, we drastically reduce time by combining all the stains into a clearing solution, applied to the sample at the same time. This is opposed to applying stains sequentially, with wash steps in between. All fixing, staining, and washing steps are completed directly on glass slides, reducing mechanical damage, ultimately making the staining process more convenient. The entire method takes about 4 hours and requires less than 500 µL of combined staining and clearing solution per slide/sample. This pipeline allows for the preparation of many slides at a time, increasing throughput without compromising anatomical and cellular features. We are able to maintain quantitative power by using three cell wall stains, Calcofluor White, Fluorol Yellow 088, and Basic Fuchsin, which are visualized at different fluorescent channels and stain cellulose, suberin, and lignin, respectively. As a proof of concept, we utilized this method for investigating the root developmental series of 3 plant species (*Triticum aestivum, Zea mays, and Cucurbita pepo*), visualizing the stems of 4 different plant species (*Cucurbita pepo, Solanum lycopersicum, Tagetes patula, and Camelina sativa*), and performing quantitative analysis using ImageJ/Fiji. We focused on investigating the developmental trajectories of embryonic roots as they are vital to whole plant development and are initially responsible for the early acquisition of water and nutrients (Du et al., 2026; Hochholdinger et al., 2018). The images produced using this staining method are compatible for quantification using the open-source image processing software ImageJ/Fiji. Overall, this preparation method can be implemented across a broad range of monocot and dicot species, with various combinations of staining dyes, to help us to advance our understanding of physiology and development at the cellular level.

## 2. Results

### Slide processing is simplified by confining all steps directly onto glass slides

Traditional staining methods are highly variable and depend on many factors including the sample type and the stains used (Supplemental Table 1). For instance, Sudan III, used to visualize suberin lamellae, is usually dissolved into ethanol/water (1:1; v/v) and the staining works well at 70°C (Ouyang et al., 2020; Spicher, 1985). In contrast, phloroglucinol/HCl staining, used for visualizing lignin, works at room temperature but requires a mixture of 10% (v/v) H_2_SO_4_ in 75% (v/v) glycerol to maintain the staining color (Jensen, 1962; Ouyang et al., 2020). The requirement of different solvents, distinct optimal temperatures, and incubation time periods makes the routine process of staining experimental samples laborious and time consuming. We circumvented this issue by dissolving the staining dyes into a single solvent, ClearSee, to stain and clear root sections simultaneously at room temperature (Figure 1g-i). Additionally, transferring the sectioned root tissue from one staining solution to the next causes mechanical injuries and can be inconvenient with delicate root tissues. To avoid such damage, we sectioned fresh roots using a vibratome and placed each section directly onto a glass slide where we performed all steps including fixing, clearing, staining, and washing (Figure 1f). To keep the sample intact and protect the stain from fading, we added 50% glycerol to the sample before sealing and storage of the slide (Figure 1n). As a consequence, the in-slide staining protocol requires less time, becomes more convenient, and produces images without any mechanical injuries.

**Figure 1:**
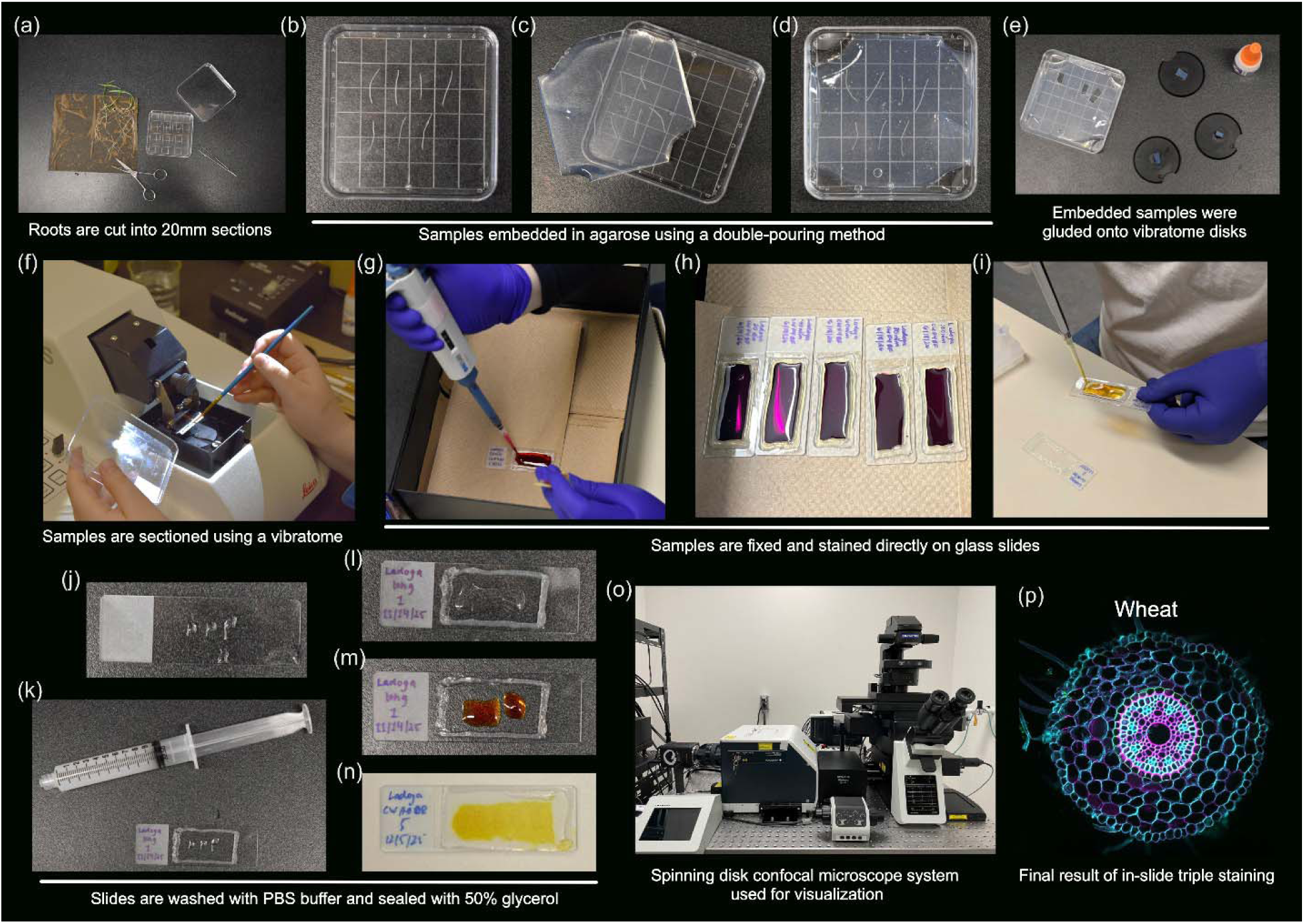
Step by step workflow of in-slide sample preparation. (a) Wheat roots grown in wet paper were cut into pieces and prepared for embedding. (b-d) Double agarose pouring method where an initial layer of agarose was poured, left to dry, and then followed up by a second layer of agarose to fully encase the samples. (e) Blocks were then cut using a razor blade and glued onto vibratome disks for sectioning. (f) Sectioned tissues were collected off of the vibratome and placed onto glass slides using a paintbrush. (g-m) Vacuum grease was piped onto glasses slides before fixing, staining, and washing samples. (n) Slides were sealed and preserved with 50% glycerol. (o) Prepared samples were imaged with a spinning disk confocal microscope. (p) Merged image of wheat root cross section depicted in figure 2.

### All staining solutions are combined into one master mix

Most investigations suggest the sequential application of stains, that is one stain applied to the sample at a time with washing steps in between (Supplemental Table 1). To increase efficiency, we decided to test whether it was possible to combine all stains into one mix and still get an effectively stained sample. Initially, we began with a combination of 2 cellulose stains (Calcofluor White and Direct Red 23) and the non-specific suberin and lignin stain Auramine O applied to image a section 6-8 cm from the root tip in both wheat and maize (Supplemental Figures 1-2). We verified the staining quality by checking for uniform fluorescence and commonly recognizable apoplastic barriers, such as the endo- and exodermis, which are known to be stained by Auramine O due to their high suberin and lignin content (Piccinini et al., 2024). Combining all the stains into one solution did not significantly impact the quality of the images. In maize, we were able to observe both uniform fluorescence from Calcofluor White and Direct Red 23 and clearly recognizable endo- and exodermis with Auramine O (Supplemental Figure 2). The same was also consistent in the case of wheat (Supplemental Figure 1). To further prove that combining different stains into one solution had no effect on image quality, we utilized a combination of Calcofluor White, Fluorol Yellow 088, and Basic Fuchsin to stain wheat (Figures 2 and 3) and maize (Figures 4 and 5) 14 cm above the root tip. These stains have previously been shown to be compatible while effectively staining distinct cell wall components (Sexauer et al., 2021). Again, we observed uniform fluorescence from all 3 stains and clearly stained endo- and exodermis in both wheat and maize (Figures 2 and 4). Thus, for the rest of our investigations, we utilized all stains dissolved into one solution applied directly on glass slides in order to accurately and quickly identify different cell wall components at different developmental zones.

**Figure 2:**
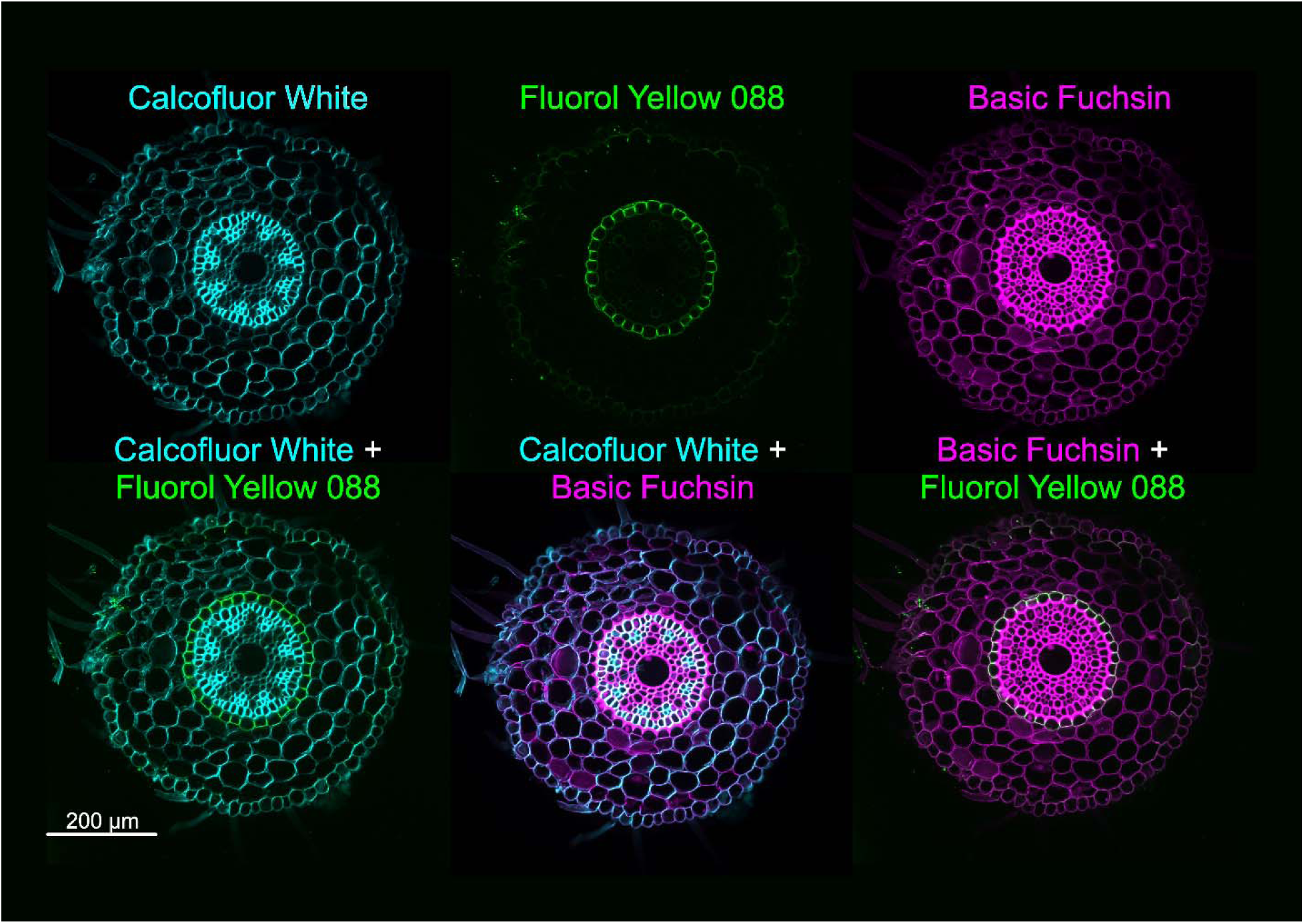
Triple stained wheat primary root cross section with Calcofluor White, Fluorol Yellow 088, and Basic Fuchsin. Wheat primary root cross section was taken from the 14 cm position or below the base of the plant. The top panels demonstrate the individual stains of Calcofluor White, Fluorol Yellow 088, and Basic Fuchsin. The bottom panels highlight ease of visualization through a combination of stains; Calcofluor White+Fluorol Yellow 088, Basic Fuchsin+Fluorol Yellow 088, and Calcofluor White+Basic Fuchsin. Scale bar = 200 µm.

**Figure 3:**
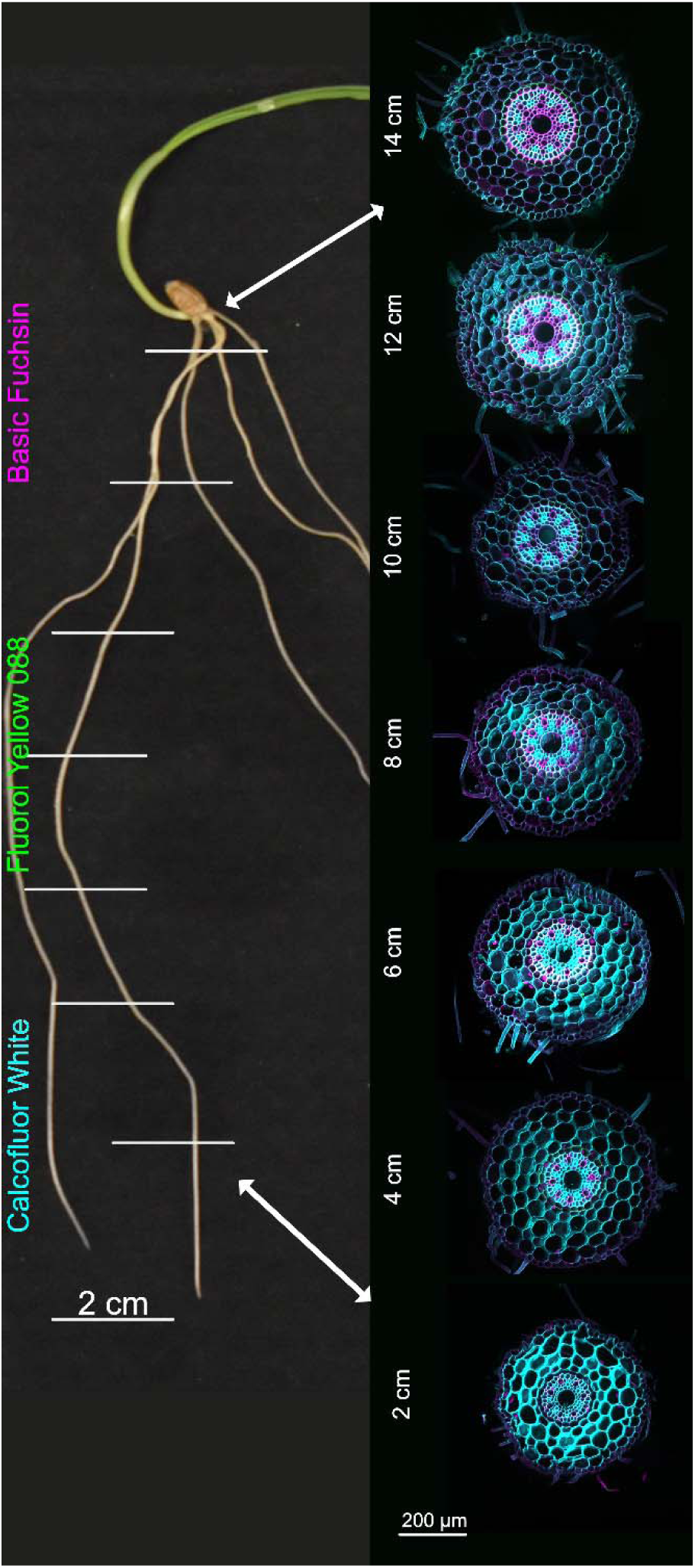
Triple stained wheat primary root developmental series with Calcofluor White, Fluorol Yellow 088, and Basic Fuchsin. Wheat primary root cross sections from 7 positions along the root (2 cm, 4 cm, 6 cm, 8 cm, 10 cm, 12 cm, and 14 cm) were taken and stained. Cross sections show internal anatomy including epidermis, exodermis, cortex, endodermis and stele. Scale bar = 200 µm.

**Figure 4:**
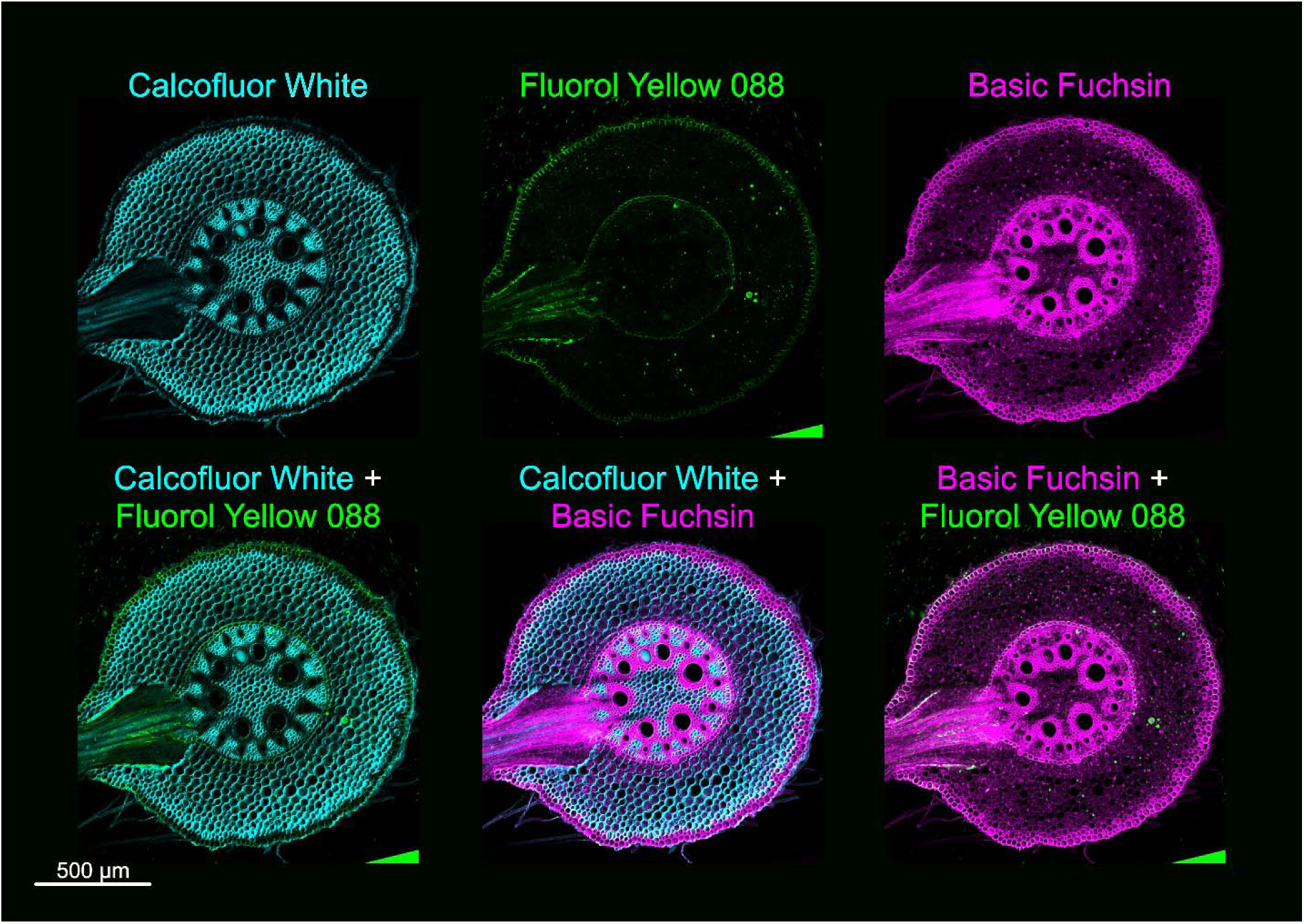
Triple stained maize primary root cross section with Calcofluor White, Fluorol Yellow 088, and Basic Fuchsin. Maize primary root cross section was taken from the 14 cm position or below the base of the plant. The top panels demonstrate the individual stains of Calcofluor White, Fluorol Yellow 088, and Basic Fuchsin. The bottom panels highlight ease of visualization through a combination of stains; Calcofluor White+Fluorol Yellow 088, Basic Fuchsin+Fluorol Yellow 088, and Calcofluor White+Basic Fuchsin. Scale bar = 500 µm.

**Figure 5:**
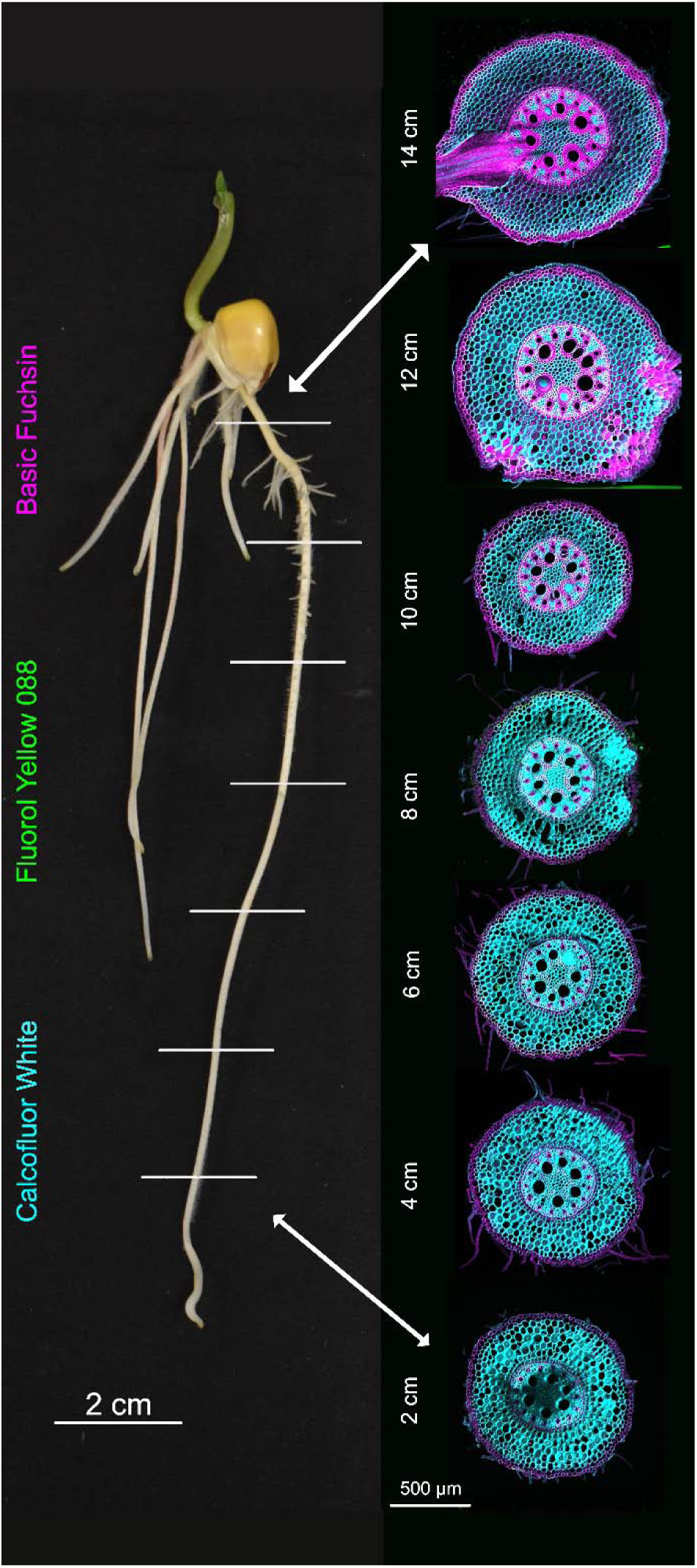
Triple stained maize primary root developmental series with Calcofluor White, Fluorol Yellow 088, and Basic Fuchsin. Maize primary root cross sections from 7 positions along the root (2 cm, 4 cm, 6 cm, 8 cm, 10 cm, 12 cm, and 14 cm) were taken and stained. Cross sections show internal anatomy including epidermis, exodermis, cortex, endodermis and stele. Scale bar = 500 µm.

### In-slide sample preparation helps to illuminate the developmental trajectory

In-slide staining enables the preparation of many samples at once and helps to illuminate the difference in cell wall composition based on developmental zone. Within our initial investigations of wheat and maize, we clearly observed root apoplastic barriers with Auramine O using in-slide sample preparation (Supplemental Figures 1 and 2). We define the outermost barrier more specifically as an exodermis, due to the prevalence of lignin and suberin deposited in the hypodermis, or the outermost cortical cell layer (Figures 2 and 4)(Liu and Kreszies, 2023). To track the progressive formation of this barrier over the course of root development, we took wheat and maize cross sections from every 2 cm beginning at the root tip and ending at the base of the plant (2 cm, 4 cm, 6 cm, 8 cm, 10 cm, 12 cm and 14 cm positions) (Figures 3 and 5, Supplemental Figures 3 and 4). We decided to use the staining combination of Calcofluor White, Fluorol Yellow 088, and Basic Fuchsin to pinpoint the specific components present in distinct developmental positions (Figures 2-5). At the 2 cm position, the most prevalent component throughout the section is cellulose, stained by Calcofluor White, with no formed exodermis, as depicted by the lack of Basic Fuchsin and Fluorol Yellow 088 staining (Figure 3 and 5). Further up the root, in addition to cellulose, lignin and suberin deposits, visualized by Basic Fuchsin and Fluorol Yellow 088, are visible in the cells that comprise the exodermis between the 4-6 cm positions (Figure 3 and 5). Finally, from the 8-14 cm positions, a clear exodermis is formed and surrounds the rest of the root tissues, as evidenced by intense fluorescence of all the stains (Figure 3 and 5). In the specific case of maize, we saw a multiseriate exodermis, or an exodermis that consists of two or more cell layers, at the 12 and 14 cm positions indicated through bright Basic Fuchsin staining (Figure 5). Ultimately, the efficiency and accuracy of in-slide sample preparation allowed us to track tissue formation and discover developmental differences in cell wall composition.

### In-slide sample preparation is convenient for use with different species and tissues

In-slide sample preparation is not only limited to use with monocot roots and can be applied to different species and tissue types. We examined the developmental trajectory of a dicot primary root to test if our preparation method worked with a different species. Similarly to wheat and maize, we took cross sections of *Cucurbita pepo* (winter squash) primary roots from every 2 cm (2 cm, 4 cm, 6 cm, 8 cm, 10 cm, 12 cm and 14 cm positions) and stained with Calcofluor White, Fluorol Yellow 088, and Basic Fuchsin (Figure 6). The cell wall composition and the formation of apoplastic barriers in winter squash are different to that of wheat and maize. In winter squash, the absence of Basic Fuchsin and Fluorol Yellow 088 staining in the hypodermis points to the lack of an exodermis at all root positions (Figure 6). Moreover, the endodermis, the inner-most apoplastic barrier, begins to accumulate lignin and suberin at the 10 cm position, exhibited through Basic Fuchsin and Fluorol Yellow 088 staining. Finally, cellulose remains the most common component through all root positions, as shown with Calcofluor White (Figure 6).

**Figure 6:**
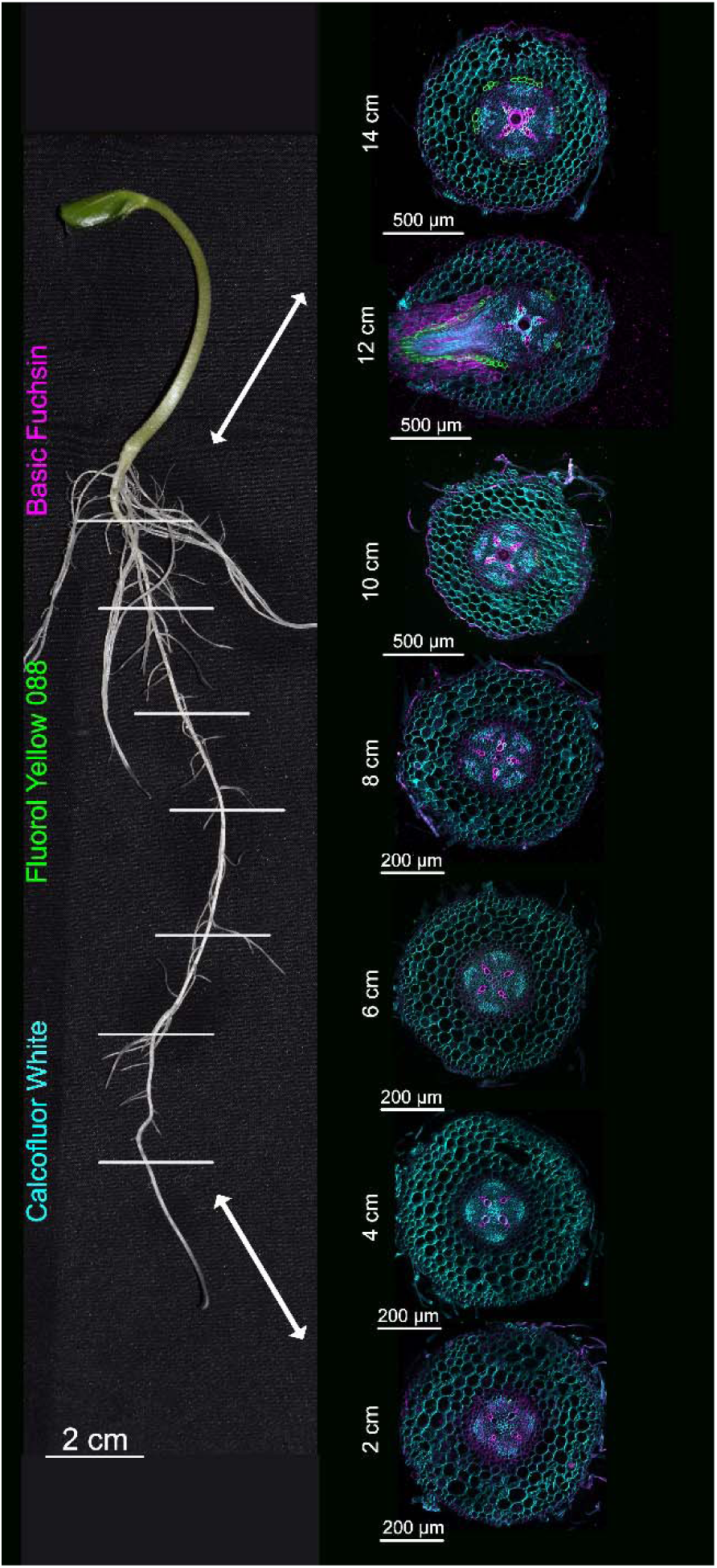
Triple stained winter squash primary root developmental series with Calcofluor White, Fluorol Yellow 088, and Basic Fuchsin. Winter squash primary root cross sections from 7 positions along the root (2 cm, 4 cm, 6 cm, 8 cm, 10 cm, 12 cm, and 14 cm) were taken and stained. Cross sections show internal anatomy including epidermis, cortex, endodermis and stele. Scale bar = 500 µm.

Furthermore, we tested the viability of the protocol with other organs like the stem (Figure 7). Stems of winter squash (*Cucurbita pepo*), tomato (*Solanum lycopersicum*), marigold (*Tagetes patula*), and false flax (*Camelina sativa*) were utilized to represent a wide range of plant lineages. All sections were easily visualized as evidenced by uniform staining and fluorescence (Figure 7). While specific staining patterns varied between different organisms, the stems were characterized by intense Basic Fuchsin staining in the middle of the sections, depictive of lignin-based vascular reinforcement (Emonet and Hay, 2022). Additionally, with the exception of small regions in the winter squash and marigold stems, there is a relative lack of Fluorol Yellow 088 stained suberin (Figure 7). Altogether, in-slide sample preparation is viable for use with dicots and different organs other than the root and can be immediately used to identify developmental differences between species.

**Figure 7:**
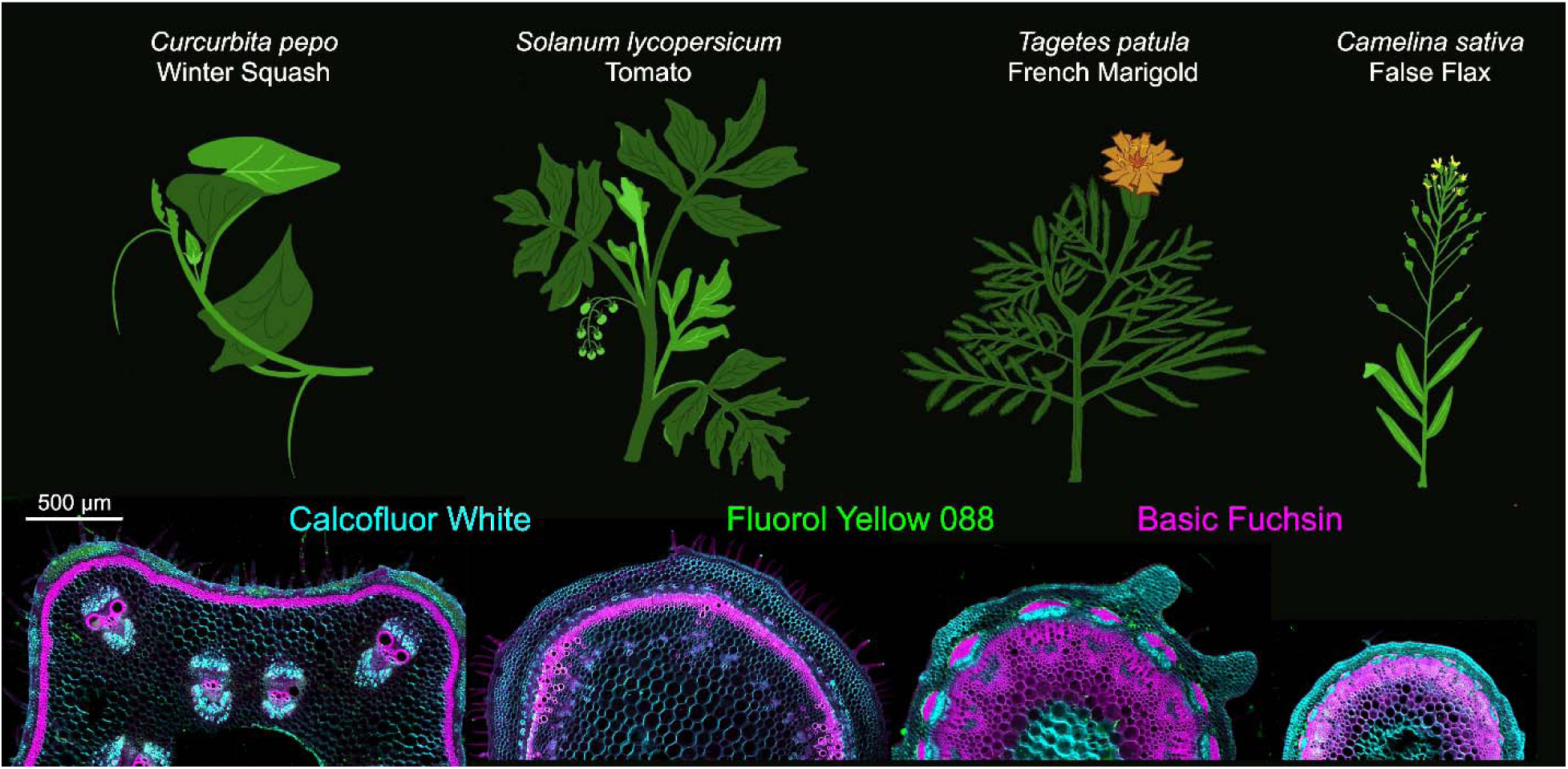
Triple stained mature stems with Calcofluor White, Fluorol Yellow 088, and Basic Fuchsin. Stems from apical growth points of winter squash, tomato, marigold, and false flax were excised, sectioned and stained. Cross sections show internal anatomy including epidermis and vasculature. Representative pictures of each species are hand drawn. Scale bar = 500 µm.

### Quantitative cell biology approach for root development

The nature of in-slide sample preparation, in which many slides are stained with the same mixture at the same time (Figure 1h), results in consistent staining across all samples. This methodological consistency further allows for comparisons based on staining fluorescence intensity, which is thought to be representative of the cell wall materials present. Here, we present 2 options; measuring spatiotemporal fluorescence at the single cell level (Figure 8) or the total fluorescence of whole sections (Supplemental Figure 5). To measure cellular spatiotemporal fluorescence, we took advantage of maize root cells with a developing exodermis at the 10 cm position (Figure 8). We drew a 70 µm line across a cell from the exodermis either in the horizontal or vertical direction to investigate whether the fluorescence varied based on location (Figure 8). Using the “plot profiler” function in ImageJ/Fiji, the intensity of Calcofluor White is relatively high compared to the other cell wall components and similar for both the horizontal and vertical directions (Figure 8). In the case of Fluorol Yellow 088, the signal intensity from horizontal direction is relatively equal. However, the intensity observed from the vertical direction is significantly lower than that of the horizontal, highlighting the presence of anticlinally deposited suberin in the horizontal direction. For Basic Fuchsin, the fluorescence is less intense compared to that of Calcofluor White, but it follows a similar pattern where the horizontal and vertical directions have similar intensities (Figure 8). Additionally, we measured the total fluorescence of the maize root sections across all 3 different channels using ImageJ/Fiji (Supplemental Figure 5). The resulting values confirm the initial observation of lower fluorescence in the regions closer to the root tip (2-4 cm positions) compared to those in the regions closer to the base (10-14 cm positions)(Supplemental Figure 5 and Figure 4). In conclusion, in-slide sample preparation facilitates quantitative cell biology approaches to study the cell wall materials of different developmental zones.

**Figure 8:**
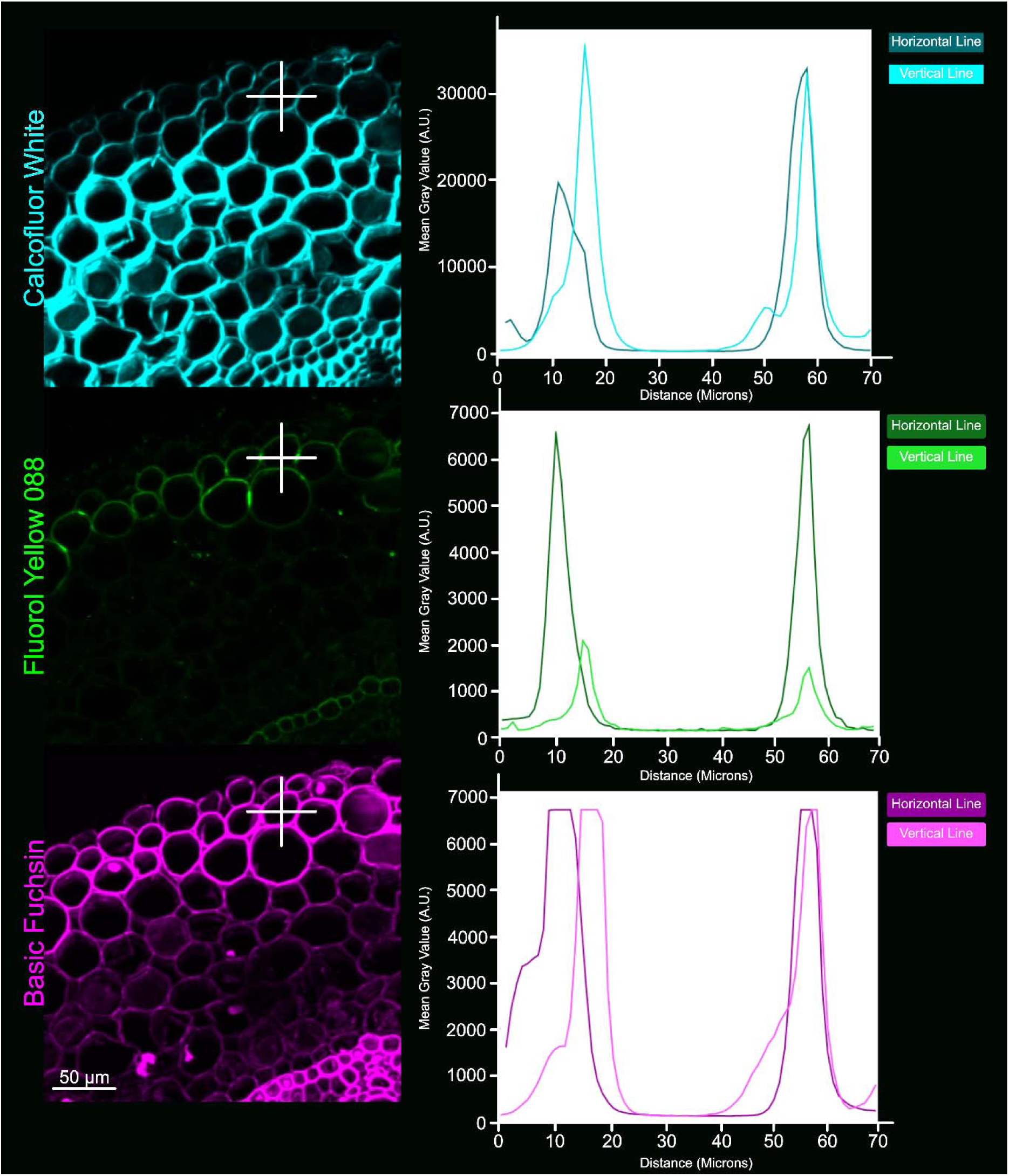
Cell type specific quantification of Calcofluor White, Fluorol Yellow 088, and Basic Fuchsin using ImageJ/Fiji. Maize root exodermis cells from the 10 cm position were used for quantification. Plot profiler highlights the intensity of Calcofluor White (top), Fluorol Yellow 088 (center), and Basic Fuchsin (bottom) from horizontal and vertical directions of the cell, depicted by the white lines. Scale bar = 50 µm.

## 4. Discussion

Overall, we have developed a time efficient, inexpensive, and robust method, in-slide sample preparation, to observe and quantify root anatomical structures and cell wall components using the open-source software ImageJ/Fiji. We have developed an easy way to fix and stain samples using a single glass slide (Figure 1). Our method saves time, fixative chemicals, and staining solutions while greatly reducing mechanical injuries. Samples produced with our method provide subsequent images that are properly stained and consistent across many different developmental regions, aiding in quantification that limits human error and bias (Figures 1 and 8). Previous studies showed the benefit of dissolving staining dyes directly into the main clearing solution, which allows tissue to be cleared and stained at the same time (Manzano et al., 2025; Sexauer et al., 2021). In our described protocol, we further utilized three stains dissolved into the same clearing solution applied to the sample at once (appropriate for imaging at the same time through 405nm, 488nm, and 561nm laser lines). Using in-slide sample preparation, we were able to track the formation of root apoplastic barriers like the exodermis and also examine differences in cell wall composition based on developmental zone (Figures 2-6). Finally, we demonstrated that this preparation method is compatible for cellular level quantification and can be used with a wide array of species and in tissues other than the root, such as the stem.

Initially, we utilized a combination of 2 cellulose stains, Calcofluor White and Direct Red 23, with Auramine O (Supplemental Figures 1 and 2). Due to the differential binding specificities of the Calcofluor White and Direct Red 23, stain competition likely occurred. This was further supported by the necessity of a higher concentration of Calcofluor White to achieve an adequate fluorescence signal compared with the other combination of stains used further on in the study (Supplemental Method 1). Subsequently, while we identify that in-slide staining is an “all in one” protocol that streamlines processing for roots and stems, an additional clearing step is best utilized for organs like the leaves as they have extremely high levels of competing fluorophores (Donaldson, 2020). Such a step can either be incorporated into the in-slide preparation pipeline or before sectioning. An interesting possibility would be to attempt to combine other stains, such as DAPI or Nile Red, into a homogeneous mix as shown here. The only limitation concerning this endeavor would be that each stain must be visualized at distinct laser lines to confidently distinguish organelles or cell wall components from each other (Piccinini et al., 2024). Nonetheless, we have shown that histochemical staining is more forgiving than previously thought.

In our investigation of root developmental trajectories, we were able to identify the formation of an exodermis in wheat under control conditions (Figure 2). The wheat exodermis is thought to be an inducible layer under many types of abiotic and biotic stresses as opposed to a constitutive tissue type (Liu and Kreszies, 2023; Ouyang et al., 2020). Prior data postulates the absence of an exodermis closest to the root tip and throughout three-quarters of the root, with an exodermis present closest to the base (Ouyang et al., 2020). Moreover, a comparative study examining apoplastic barrier physiology in hydroponically and soil grown roots identified exodermis formation in maize only and not in wheat (Carvajal et al., 2025). However, our method not only detects the presence of an exodermis, but also the gradual formation of it starting as early as 4 cm from the wheat root tip (Figure 3). Additionally, this may be the first report of an exodermis prevalent in wheat embryonic roots and of a multiseriate exodermis in maize embryonic roots. A possible explanation is that “Ladoga”, the wheat variety used in this particular study, has an easily detectable or highly sensitive exodermis barrier. Monocots, especially wheat, display immense phenotypic and varietal diversity showcased both by whole root system and anatomical investigations. For instance, a study of 184 wheat genotypes from different countries revealed significant differences in root length(Chen et al., 2020). Another study on multiseriate cortical sclerenchyma (MCS) development in maize showcased a group of wheat varieties with individuals that had or completely lacked MCS (Schneider et al., 2021). Such early detection of shifting developmental components emphasizes the necessity for increased investigation of wheat embryonic and post-embryonic roots at the cellular level and has vast implications as a diagnostic tool.

An important application of this protocol could be for the screening of cell type specific developmental root traits in a variety of germplasm. It has been proposed that modern monocot root systems are inadvertently fluctuating towards reduced hydraulic capacity in favor of manageable resiliency compatible with efficient agriculture (Baca Cabrera et al., 2025; Rinehart et al., 2024). To combat this, an effort to investigate hidden and helpful root traits at both the phenotypic and genetic levels must be undertaken. In the past, root system phenotyping was unapproachable or did not tell a complete story of root development (Atkinson et al., 2019). Common soil-based root phenotyping methods, such as shovelomics (Arifuzzaman et al., 2019; Voss-Fels et al., 2018) or the use of X-ray CT scanners (Teramoto et al., 2020), are great for visualizing whole system qualities such as root system architecture. Whereas early developmental investigations, aided by hydroponics (Zeng et al., 2024) or rolled paper towel-based methods (Draves et al., 2022), provide key insight into unique root traits otherwise not captured by soil-based methods, such as altered root hair phenotypes (Zheku et al., 2026). Further efforts, such as in-slide sample preparation, to enable root phenotyping at the cellular level, can help to directly identify cell specific physiological traits and response to climate change (Hazelwood et al., 2026). Our method can enhance the capability of low-cost root anatomical phenotyping platforms, such as the Rapid Anatomics Tool (RAT)(Jones et al., 2026). Altogether, our method, in the context of other traditional root phenotyping methods, provides a new way to investigate root developmental zones and spatiotemporal distribution of cellular components with the goal of improving anatomical traits in a challenging climate.

## 5. Materials and Methods

### Plant Materials and growth conditions

Wheat (variety “Ladoga,” Prairie Garden Seeds; https://prairiegardenseeds.ca/), maize (inbred line B73 from MaizeGDB; https://www.maizegdb.org/), and winter squash (variety “Table Queen,” West Coast seeds; https://www.westcoastseeds.com/) were initially germinated on square petri dishes (Cat #26-275; Genesee Scientific) with a moist paper towel. Wheat and maize seeds were placed on plates lined with a moist paper towel, covered with aluminum foil and vernalized for two days at 4. Afterwards, the dishes were placed horizontally on a shelf at 23 with a 24 h light cycle for the plants to germinate and grow for two days. Winter Squash seeds were placed directly on the growth rack without vernalization until the seeds cracked open and germinated (usually about 2 days.) The germinated seedlings were then transferred to transparent boxes (16.5 × 8.5 cm; Temu; https://www.temu.com/) also lined with a moist paper towel. The seedlings were left to grow until the primary root measured about 14 cm in length (a total of four days for Wheat and maize or a total of three days for winter Squash) at 23 with a 24 h light cycle.

Stems of Marigold (variety “Sparky Mix,” West Coast seeds; https://www.westcoastseeds.com/), false flax (*Camelina sativa*, unknown variety) winter squash, and tomato (variety “M82”) were also utilized to verify the protocol. The plants were grown in the greenhouse with an ambient temperature range of 25-35 and a 16 h light/8 h dark cycle until mature development (2 to 3 months). Stems were excised from apical growth points of the plants and stored in 100% ethanol (Commercial alcohols Inc, https://greenfield.com/commercialalcohols/) before sectioning the same day.

### Fixation liquid, stain and ClearSee preparation

To prepare the fixative solution, 10 mL of 37% Formaldehyde (Cas no. 50-00-0, Fisher Scientific, https://www.fishersci.com/us/en/home.html) was added to 50 mL of 100% ethanol and 40 mL of sterilized water to produce 100 mL of 4% formaldehyde. Throughout the development of this protocol, the fixative solution was reused with no issue in sample fixing or staining. Thus, it is not necessary to use fresh fixative for this protocol as long as the solution is kept clean of debris and cross contamination (Supplemental method 1). Initial stock solutions and working volumes of the stains were prepared with ClearSee (Kurihara et al., 2015). Fluorol Yellow 088 stock solution was prepared as per reference and then incorporated into the ClearSee based mastermix (Sexauer et al., 2021). This method was designed to allow for staining and clearing of tissue to occur during the same step. ClearSee was prepared using the protocol described by Kurihara et al., 2015 (Supplemental method 1). The staining chemicals used include 5 mM Calcofluor White (Cas no. 4404-43-7, bioWORLD, https://www.bio-world.com/), Auramine O (Cas no. 2465-27-2, Sigma Aldrich, https://www.sigmaaldrich.com/CA/en), Direct Red 23 (Cas no. 3441-14-3, Sigma Aldrich, https://www.sigmaaldrich.com/CA/en), Fluorol Yellow 088 (ChemCruz, Cas no.81-37-8, https://www.scbt.com/p/fluorol-yellow-088-81-37-8), and Basic Fuchsin (Sigma Aldrich, Cas no.569-61-9, https://www.sigmaaldrich.com/CA/en). The stain master mix was prepared to reach working volumes of 0.0625 mM Calcofluor White, 0.01% (w/v) Fluorol Yellow 088, and 0.2% (w/v) Basic Fuchsin in 1 mL of ClearSee (Supplemental method 1). Alternatively, another mix of 0.25mM Calcofluor white, 0.5% (w/v) Auramine O, and 0.1% (w/v) Direct Red 23 in 1 mL of ClearSee can also be utilized (Supplemental Figure 1-2). Combining all stains into one master mix saves time spent staining as well as the amount of total stain used. For example, on a typical day, the whole process can take up to 4 hours, depending on the number of slides made, and uses only about 1 mL of prepared stain. A new stain master mix was prepared fresh at the time of sectioning, covered with tinfoil, and kept at room temperature until use.

### Block preparation and sectioning

Block preparation and sectioning protocols are consistent for all plant parts. Sectioning blocks were prepared, using square petri dishes as molds, by submerging root and stem samples in 5% agarose (Genesee Scientific, Cas no. 90120-36-6, https://www.geneseesci.com/). A double pouring method was developed to ensure correct orientation of samples, which helps immensely in obtaining clean sections later (Figure 1b-d). First, an initial layer of agarose was poured, and samples were quickly placed inside using forceps, taking care to ensure correct alignment in the block, and left to solidify completely. Then, the first layer of agarose was gently removed from the mold. Another thin layer of agarose was poured into the mold, with the first layer containing the samples gently placed on top afterwards. This way, the sample is embedded fully in the agarose, with thin layers protecting it on each side. Using a razor blade, the block was cut into manageable pieces for use with the vibratome (Leica VT1000 S; https://www.leicabiosystems.com/) (Figure 1e). Blocks were glued onto vibratome disks with cyanoacrylate glue and placed onto the machine with sterilized water for sectioning. Cross sections were 100 µM thick. Sections coming off of the vibratome were immediately picked up with a paintbrush and organized onto slides (Figure 1f). Quality of the sectioned sample was verified with a brightfield microscope (AmScope T490 Series Simul-Focal Biological Trinocular Compound Microscope; https://amscope.com/) before staining (Supp. Figure 3-4).

### In-slide preparation of sectioned sample

All staining steps took place at room temperature. After verification of the sample with brightfield microscopy, the slide was labelled accordingly and excess water removed with a kimwipe. A thin layer of vacuum grease was piped using a syringe around the sample, taking care to not add too much or be too close to the sample. The vacuum grease acts as a makeshift well for the staining steps later on. Extra care was taken to make sure the slide was completely dry before adding the vacuum grease, as good contact of the vacuum grease with the slide helps to avoid spills later on. For each step, enough liquid was added to the vacuum grease well to completely cover the sample and fill the well. First, the sample was fixed with 4% fixative solution for 30 minutes. Excess fixative was pipetted off the slide and reused. Then, stain master mix was added and the sample stained for a total of 3 hours, with gentle agitation every 15 minutes. Excess stain was washed off the slide with 1× PBS (Thermo Fisher Scientific, https://www.thermofisher.com/ca/en/home.html) 3 to 4 times. Finally, samples were reoriented on the slide using a paintbrush before adding 50% glycerol (Fisher Scientific, Cas no. 56-81-5, https://www.fishersci.ca/ca/en/home.html) and a cover slip on top of the vacuum grease to finish. Slides were stored in the dark at 4 and can be used up to two to three weeks after preparation.

### Confocal microscopy

Confocal images were taken using Evident IXplore SpinSR Spinning Disk Confocal System at the Bioimaging Facility (BIF, https://www.bioimaging.ubc.ca/equipment/lightmicroscopes/evident-ixplore-spinsr-spinning-disk-confocal-system/) of the University of British Columbia (UBC), Vancouver, Canada. The confocal system is equipped with Evident Inverted IX83 microscope base, Yokogawa CSU-W1 (2 discs: 50 μm pinhole disk for high resolution confocal imaging), Marzhauzer Piezo stage, 2× Hamamatsu ORCA-Fusion BT back-thinned Gen-III sCMOS (2304 × 2304 pixel, 6.5 μm × 6.5 μm pixel size, up to 31.6 fps at 2304 × 2304 pixel). X-Cite mini+ LED for wide-field fluorescence was used to locate the sample in the slide and primary focus; and 405nm (50mW), 488nm (100mW), and 561nm (100mW) laser lines were used for Calcofluor White, Fluorol Yellow 088, and Basic Fuchsin respectively. Images were captured using 10× (UPLXAPO 10×/0.40NA, Dry), 20× (UPLXAPO 20×/0.80NA, Dry), and 40× (UPLSAPO 40×/0.95NA, Dry) objective lenses. CellSens Dimension software was used for initial acquisition and later on post processing was performed using ImageJ/Fiji (Schindelin et al., 2012).

### Image processing

Raw image data was processed using ImageJ/Fiji (Schindelin et al., 2012). To calculate fluorescence intensity, the area of interest was first selected using the segmented line tool. Fluorescence intensity was then calculated by dividing the mean grey value of the relative pixels present by the selected area. The intensity data was collected using a selection tool of 70 μm straight line and Plot Profile (ImageJ/Fiji > Analyze > Plot Profile) plugin. The intensity data obtained from ImageJ and the Plot Profile was used to make the graph using JMP Pro 17 (https://www.jmp.com/en/home).

## Supporting information

Supplemental protocol

## Acknowledgements

Authors are grateful to Dr. Miki Fujita, Dr. EunKyoung Lee, and UBC Bioimaging facility (RRID: SCR_021304) for their help and support in confocal imaging. Authors also thank Scottie Mello, Melina Biron and the UBC greenhouse technicians for their help with plant maintenance.

## Authors’ contributions

J.Z. performed experiments, developed methods, wrote the initial draft of the manuscript. J.Z., R.S., and M.A.A. together conceived the idea of the protocol and wrote the final version of the manuscript. J.Z. and M.A.A. prepared figures and performed the quantification. M.A.A. and R.S. supervised J.Z. during the project.

---

**Supplemental Figure 1:**
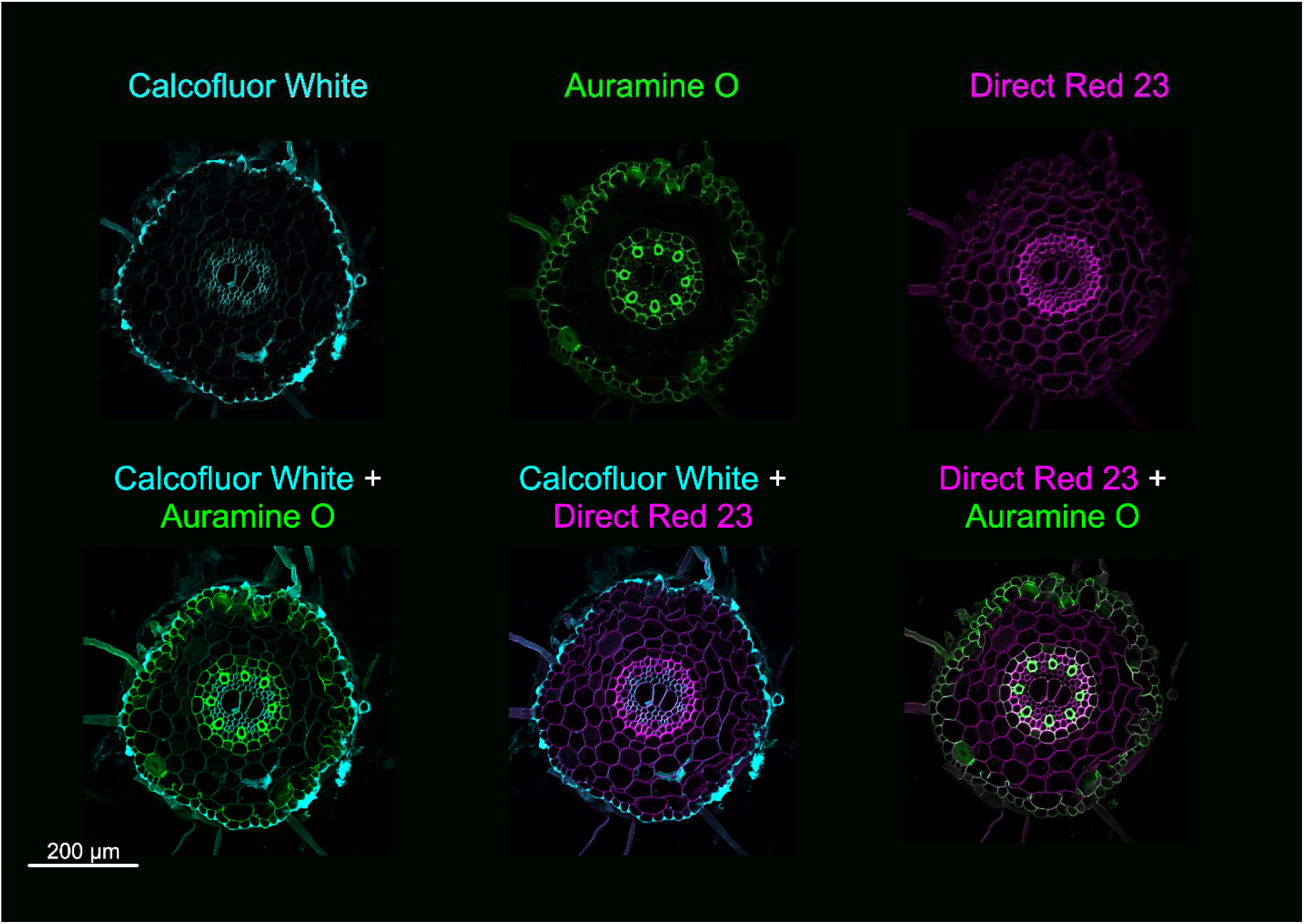
Triple stained wheat primary root cross section with Calcofluor White, Auramine O and Direct Red 23. Wheat primary root cross section was taken from the 6-8 cm position of the plant. The top panels demonstrate the individual stains of Calcofluor White, Auramine O and Direct Red 23. The bottom panels highlight ease of visualization through a combination of stains; Calcofluor White+Auramine O, Direct Red 23+Auramine O, and Calcofluor White+Direct Red 23. Scale bar = 200 µm.

**Supplemental Figure 2:**
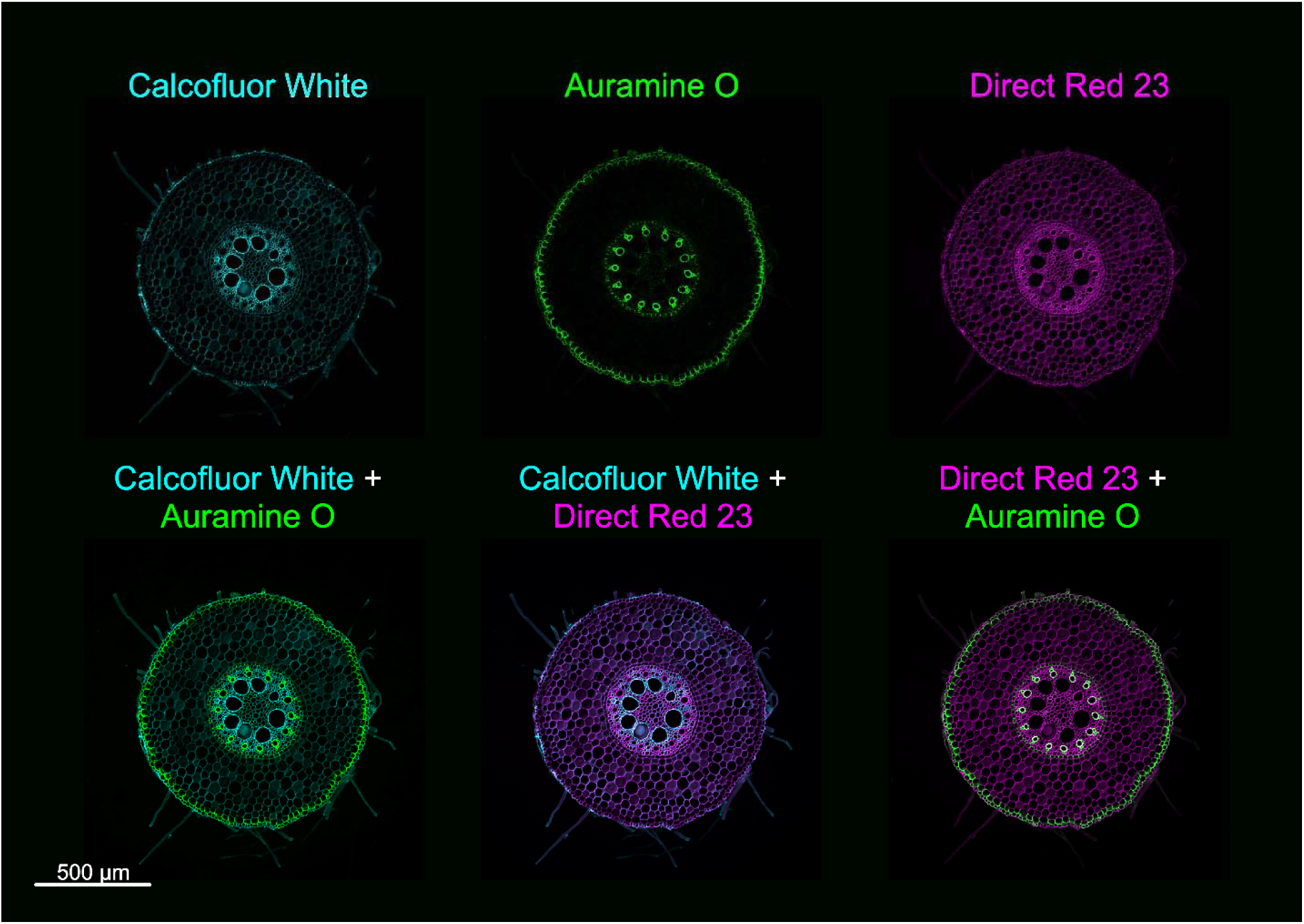
Triple stained maize primary root cross section with Calcofluor White, Auramine O and Direct Red 23. Maize primary root cross section was taken from the 6-8 cm position of the plant. The top panels demonstrate the individual stains of Calcofluor White, Auramine O and Direct Red 23. The bottom panels highlight ease of visualization through a combination of stains; Calcofluor White+Auramine O, Direct Red 23+Auramine O, and Calcofluor White+Direct Red 23. Scale bar = 200 µm.

**Supplemental Figure 3:**
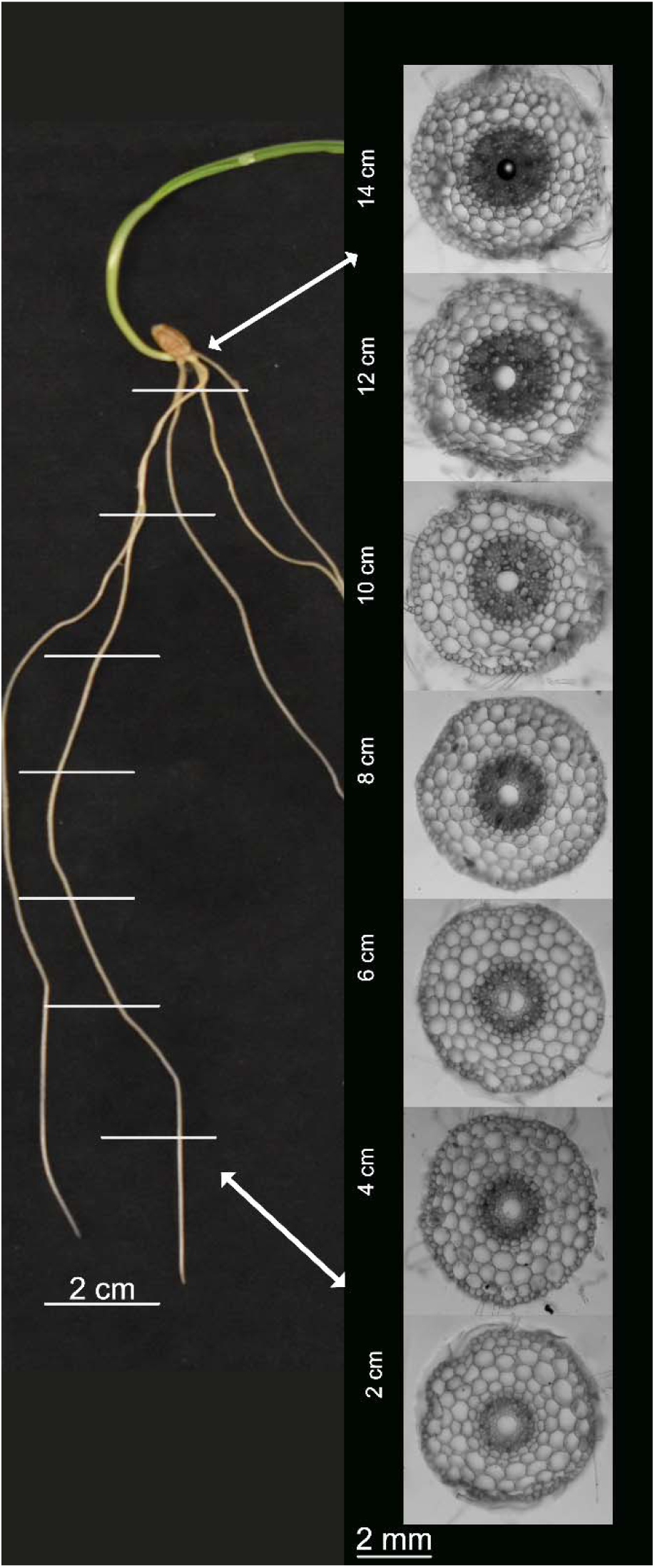
Unstained wheat primary root developmental series. Wheat primary root cross sections from 7 positions along the root (2 cm, 4 cm, 6 cm, 8cm, 10 cm, 12 cm, and 14 cm) were taken and visualized with a brightfield microscope at 10× objective. Scale bar = 2mm

**Supplemental Figure 4:**
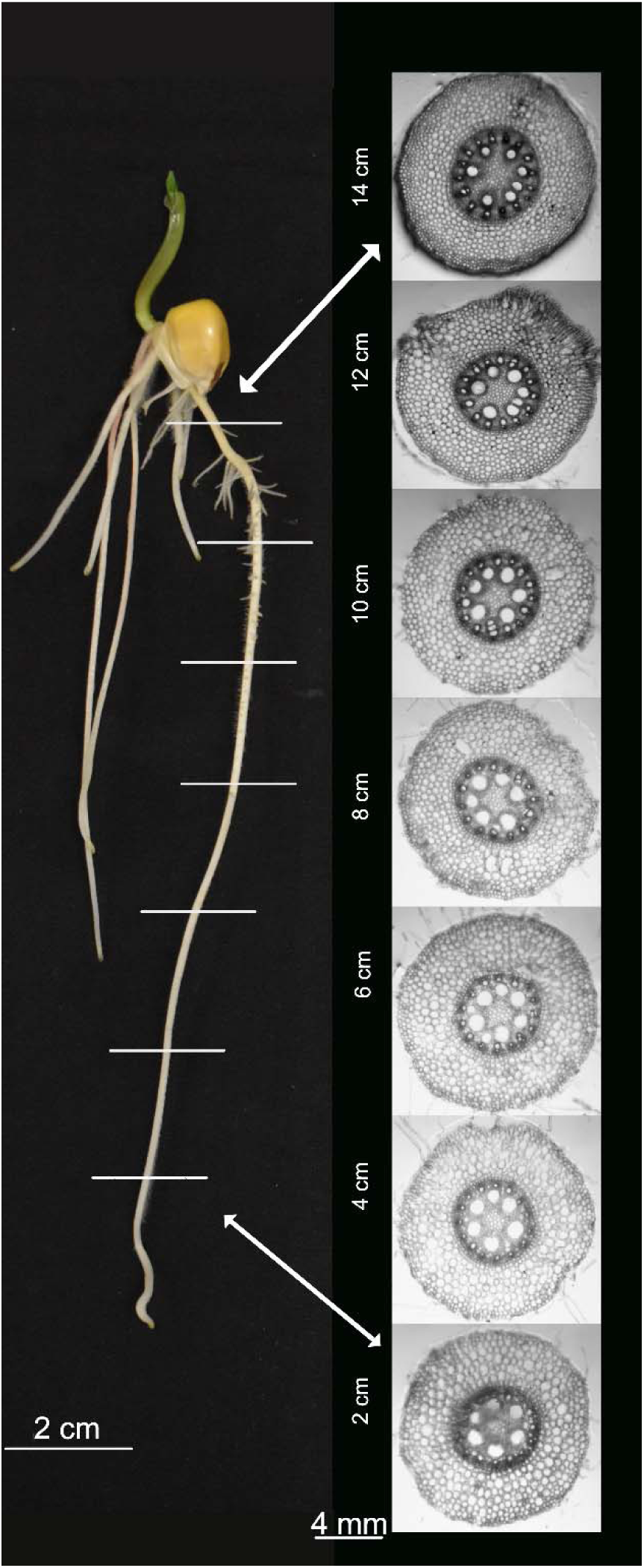
Unstained maize primary root developmental series. Maize primary root cross sections from 7 positions along the root (2 cm, 4 cm, 6 cm, 8cm, 10 cm, 12 cm, and 14 cm) were taken and visualized with a brightfield microscope at 10× objective. Scale bar = 4mm

**Supplemental Figure 5:**
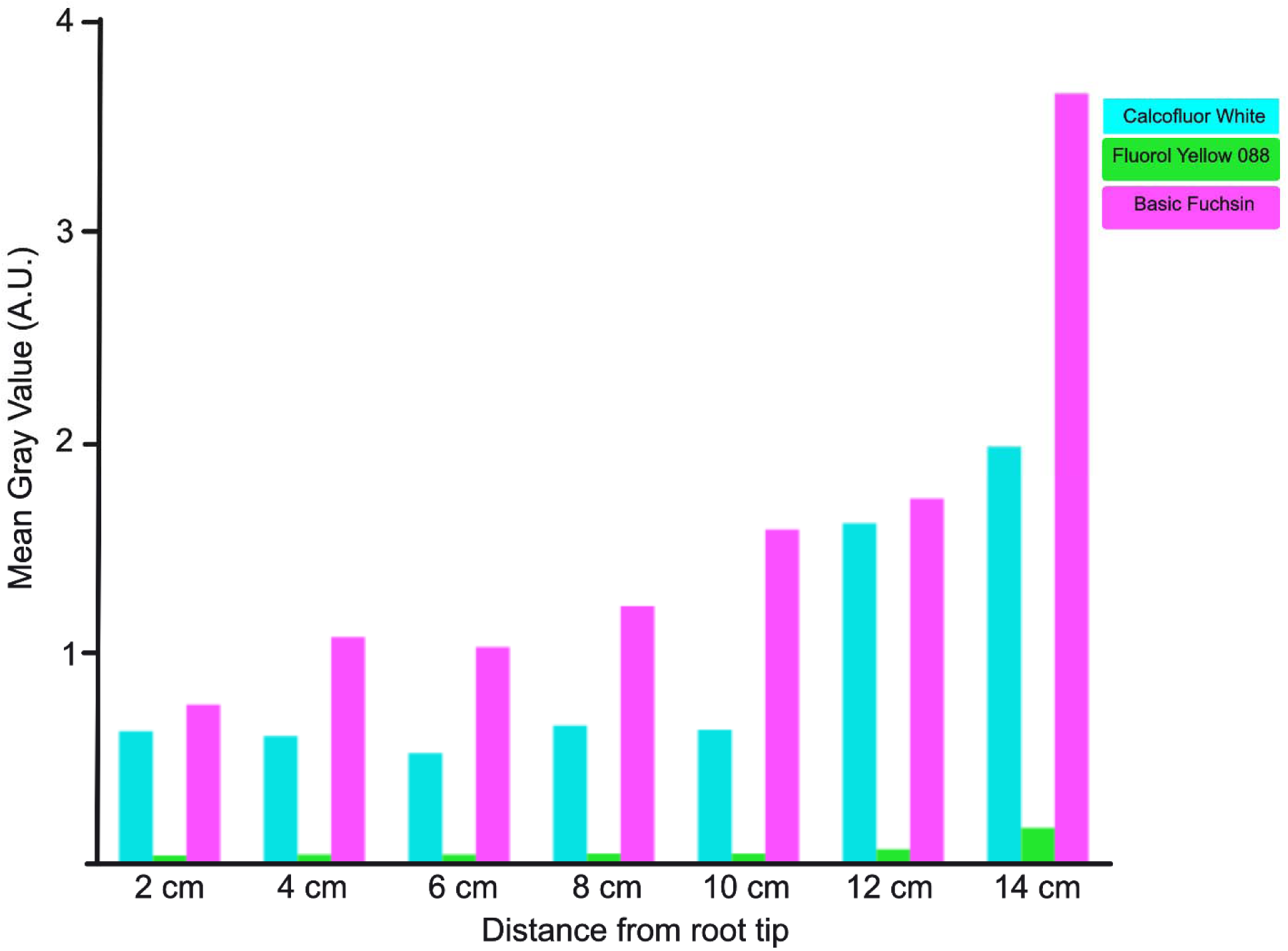
Example graph of total fluorescence intensity across different developmental zones of the maize root. Fluorescence intensity was measured using ImageJ/Fiji for each laser line that corresponded to specific cell wall components present at that developmental position (405nm for cellulose stained by Calcofluor White, 488nm for suberin stained by Fluorol Yellow 088, and 561nm for lignin stained by basic Fuchsin).

## References

Arifuzzaman, M., Oladzadabbasabadi, A., McClean, P. and Rahman, M. (2019). Shovelomics for phenotyping root architectural traits of rapeseed/canola (Brassica napus L.) and genome-wide association mapping. Mol Genet Genomics 294, 985–1000.

Atkinson, J. A., Pound, M. P., Bennett, M. J. and Wells, D. M. (2019). Uncovering the hidden half of plants using new advances in root phenotyping. Current Opinion in Biotechnology 55, 1–8.

Baca Cabrera, J. C., Vanderborght, J., Behrend, D., Gaiser, T., Boursiac, Y., Nguyen, T. H. and Lobet, G. (2025). Decreased root hydraulic traits in German winter wheat cultivars over 100 years of breeding.

Baca Cabrera, J. C., Jones, D. H., Vanderborght, J., Behrend, D., Schneider, H. M. and Lobet, G. (2026). Root anatomical gradients and cultivar differences underlie variation in root hydraulic properties in German winter wheat. J Exp Bot erag177.

Cantó-Pastor, A., Kajala, K., Shaar-Moshe, L., Manzano, C., Timilsena, P., De Bellis, D., Gray, S., Holbein, J., Yang, H., Mohammad, S., et al. (2024). A suberized exodermis is required for tomato drought tolerance. Nat. Plants 10, 118–130.

Carvajal, J., Suresh, K., Bhattacharyya, S., Zeisler-Diehl, V. V., Wojciechowski, T. and Schreiber, L. (2025). Comparing apoplastic root barrier formation and morphology in six crop species cultivated in soil vs. hydroponics. Planta 262, 141.

Chen, Y., Palta, J., Prasad, P. V. V. and Siddique, K. H. M. (2020). Phenotypic variability in bread wheat root systems at the early vegetative stage. BMC Plant Biol 20, 185.

Chen, A., Liu, T., Wang, Z. and Chen, X. (2022). Plant root suberin: A layer of defence against biotic and abiotic stresses. Front. Plant Sci. 13, 1056008.

Czymmek, K. J., Benitez-Alfonso, Y., Burch-Smith, T. and Di Costanzo, L. (2025). Best practices in plant fluorescence imaging and reporting: A primer.

Donaldson, L. (2020). Autofluorescence in Plants. Molecules 25, 2393.

Draves, M. A., Muench, R. L., Lang, M. G. and Kelley, D. R. (2022). Maize Seedling Growth and Hormone Response Assays Using the Rolled Towel Method. Current Protocols 2, e562.

Du, X., Klaus, A., Alba, M. A. G., Stöcker, T., Carvajal, J., Zeisler-Diehl, V., Schreiber, L., The, J., Consonni, G., Schoof, H., et al. (2026). dizzy1 coordinates shoot and root development, constitutive stress responses, and phytohormone signaling in maize. Front. Plant Sci. 17, 1927522.

Emonet, A. and Hay, A. (2022). Development and diversity of lignin patterns. Plant Physiol 190, 31–43.

Enstone, D. E. and Peterson, C. A. (2005). Suberin lamella development in maize seedling roots grown in aerated and stagnant conditions. Plant Cell & Environment 28, 444–455.

Hazelwood, O. S., Diehl, K. A., Hollenbeck, V., Demura-Devore, J., Herb, D., Gallagher, J. P. and Ashraf, M. A. (2026). Cell cycle follows ‘pause and play’ mechanism in salt and cold stress recovery in diverse plant species. New Phytologist 250, 1689–1703.

Hochholdinger, F., Marcon, C., Baldauf, J. A., Yu, P. and Frey, F. P. (2018). Proteomics of Maize Root Development. Front. Plant Sci. 9, 143.

Jensen, W. A. (1962). Botanical histochemistry: principles and practice. San Francisco, W. H. Freeman.

Jones, D. H., Baca Cabrera, J. C., Behrend, D., Wells, D. M., Swift, J. F., Atkinson, J. A., Schön, M., Lobet, G., Hanlon, M. T. and Schneider, H. M. (2026). The Rapid Anatomics Tool (RAT): A low-cost root anatomical phenotyping platform reveals changes in root anatomy along the root axis. Plant Phenomics 8, 100150.

Kitin, P., Nakaba, S., Hunt, C. G., Lim, S. and Funada, R. (2020). Direct fluorescence imaging of lignocellulosic and suberized cell walls in roots and stems. AoB PLANTS 12, plaa032.

Kreszies, T., Schreiber, L. and Ranathunge, K. (2018). Suberized transport barriers in *Arabidopsis*, barley and rice roots: From the model plant to crop species. Journal of Plant Physiology 227, 75–83.

Kurihara, D., Mizuta, Y., Sato, Y. and Higashiyama, T. (2015). ClearSee: a rapid optical clearing reagent for whole-plant fluorescence imaging. Development 142, 4168–4179.

Li, J., Minami, S., Tsuda, K. and Yamauchi, T. (2026). SCARECROW functions in root radial patterning in rice to organize multiple cortical cell layers.

Liu, T. and Kreszies, T. (2023). The exodermis: A forgotten but promising apoplastic barrier. Journal of Plant Physiology 290, 154118.

Manzano, C., Morimoto, K. W., Shaar-Moshe, L., Mason, G. A., Cantó-Pastor, A., Gouran, M., De Bellis, D., Ursache, R., Kajala, K., Sinha, N., et al. (2025). Regulation and function of a polarly localized lignin barrier in the exodermis. Nat. Plants 11, 118–130.

Ortiz-Ramírez, C., Guillotin, B., Xu, X., Rahni, R., Zhang, S., Yan, Z., Coqueiro Dias Araujo, P., Demesa-Arevalo, E., Lee, L., Van Eck, J., et al. (2021). Ground tissue circuitry regulates organ complexity in maize and *Setaria*. Science 374, 1247–1252.

Ouyang, W., Yin, X., Yang, J. and Struik, P. C. (2020). Comparisons with wheat reveal root anatomical and histochemical constraints of rice under water-deficit stress. Plant Soil 452, 547–568.

Piccinini, L., Nirina Ramamonjy, F. and Ursache, R. (2024). Imaging plant cell walls using fluorescent stains: The beauty is in the details. Journal of Microscopy 295, 102–120.

Rinehart, B., Borras, L., Salmeron, M., McNear, D. H. and Poffenbarger, H. (2024). Commercial maize hybrids have smaller root systems after 80 Years of breeding. Rhizosphere 30, 100915.

Roeder, A. H. K., Bent, A., Lovell, J. T., McKay, J. K., Bravo, A., Medina-Jimenez, K., Morimoto, K. W., Brady, S. M., Hua, L., Hibberd, J. M., et al. (2025). Lost in translation: What we have learned from attributes that do not translate from Arabidopsis to other plants. Plant Cell 37, koaf036.

Schindelin, J., Arganda-Carreras, I., Frise, E., Kaynig, V., Longair, M., Pietzsch, T., Preibisch, S., Rueden, C., Saalfeld, S., Schmid, B., et al. (2012). Fiji: an open-source platform for biological-image analysis. Nat Methods 9, 676–682.

Schneider, H. M., Strock, C. F., Hanlon, M. T., Vanhees, D. J., Perkins, A. C., Ajmera, I. B., Sidhu, J. S., Mooney, S. J., Brown, K. M. and Lynch, J. P. (2021). Multiseriate cortical sclerenchyma enhance root penetration in compacted soils. Proceedings of the National Academy of Sciences 118, e2012087118.

Sexauer, M., Shen, D., Schön, M., Andersen, T. G. and Markmann, K. (2021). Visualizing polymeric components that define distinct root barriers across plant lineages. Development 148, dev199820.

Spicher, G. (1985). Gerlach, D.: Botanische Mikrotechnik – Eine Einführung. 3., unveränderte Auflage. Georg Thieme Verlag, Stuttgart 1984. 311 Seiten, mit 45 Abb., flexibles Taschenbuch DM 26,80. Starch - Stärke 37, 34–34.

Teramoto, S., Takayasu, S., Kitomi, Y., Arai-Sanoh, Y., Tanabata, T. and Uga, Y. (2020). High-throughput three-dimensional visualization of root system architecture of rice using X-ray computed tomography. Plant Methods 16, 66.

Truernit, E., Bauby, H., Dubreucq, B., Grandjean, O., Runions, J., Barthélémy, J. and Palauqui, J.-C. (2008). High-Resolution Whole-Mount Imaging of Three-Dimensional Tissue Organization and Gene Expression Enables the Study of Phloem Development and Structure in *Arabidopsis*. The Plant Cell 20, 1494–1503.

Tylová, E., Pecková, E., Blascheová, Z. and Soukup, A. (2017). Casparian bands and suberin lamellae in exodermis of lateral roots: an important trait of roots system response to abiotic stress factors. Ann Bot 120, 71–85.

Ursache, R., Andersen, T. G., Marhavý, P. and Geldner, N. (2018). A protocol for combining fluorescent proteins with histological stains for diverse cell wall components. The Plant Journal 93, 399–412.

Verhertbruggen, Y., Walker, J. L., Guillon, F. and Scheller, H. V. (2017). A Comparative Study of Sample Preparation for Staining and Immunodetection of Plant Cell Walls by Light Microscopy. Front. Plant Sci. 8,.

Viana, W., Scharwies, J. D. and Dinneny, J. R. (2022). Deconstructing the root system of grasses through an exploration of development, anatomy and function. Plant Cell & Environment 45, 602–619.

Voss-Fels, K. P., Robinson, H., Mudge, S. R., Richard, C., Newman, S., Wittkop, B., Stahl, A., Friedt, W., Frisch, M., Gabur, I., et al. (2018). VERNALIZATION1 Modulates Root System Architecture in Wheat and Barley. Molecular Plant 11, 226–229.

Zeng, D., Ford, B., Doležel, J., Karafiátová, M., Hayden, M. J., Rathjen, T. M., George, T. S., Brown, L. K., Ryan, P. R., Pettolino, F. A., et al. (2024). A conditional mutation in a wheat (Triticum aestivum L.) gene regulating root morphology. Theor Appl Genet 137, 48.

Zhang, J., Liu, Z., Farrar, E. J., Li, M., Lu, H., Qu, Z., Chara, O., Mitsuda, N., Sakamoto, S., Xue, F., et al. (2025). Ethylene modulates cell wall mechanics for root responses to compaction. Nature 649, 685–692.

Zheku, J., Do, T. and Ashraf, M. A. (2026). Toxic metals increase root hair density by reducing epidermal cell length. 2026.02.03.703618.

