## Supplemental protocol for "In-slide sample preparation allows for rapid triple staining and the detection of specific cell wall components in monocot roots"

ORCIDs:
J.Z. (0009-0009-8091-5043)
R.S. (0000-0002-9345-9640)
M.A.A. (0000-0002-0879-5835)

Supplemental method 1: The STAR protocol

**Fixation liquid preparation**

Materials

37% Formaldehyde (Cas no. 50-00-0, Fisher Scientific, <https://www.fishersci.com/shop/products/formaldehyde-37-by-weight-molecular-biology-fisher-bioreagents-2/BP53125#?keyword=>)

100% Ethanol (Commercial alcohols Inc, <https://greenfield.com/commercialalcohols/>)

Millipore sterilized water 

Procedure

1. Add 10 mL of 37% formaldehyde to 50 mL of 100% ethanol
2. Bring up to volume with 40 mL of sterilized water to produce a final volume of 100 mL of 4% formaldehyde

IMPORTANT TIPS

- Fixative can be stored either on the bench or at 4℃ for long term storage
- It is acceptable to reuse fixative as long it is kept clean of debris and not discolored (while developing this method, fixative was reused with no issue in preparation and visualization of the sample)

**ClearSee preparation**

Materials

Urea (Cas no. 57-13-6, Sigma Aldrich, <https://www.sigmaaldrich.com/CA/en>)

Xylitol (Cas no. 87-99-0, Sigma Aldrich, <https://www.sigmaaldrich.com/CA/en>)

Sodium deoxycholate (Cas no. 302-95-4, Sigma Aldrich, <https://www.sigmaaldrich.com/CA/en>)

Sterilized water

Procedure

1. Gently heat 100 mL of sterilized water on a stir plate
2. Add 10 g of xylitol in batches, wait for the first batch to dissolve before adding the rest of the xylitol
3. Add 15 g of sodium deoxycholate while stirring the solution
4. Finish by adding 25 g of urea while stirring, let the solution stir for an additional 30 minutes until all components are dissolved

IMPORTANT TIPS

- ClearSee was stored on the bench throughout the development of this method

**Stain preparation**

*Stock solution preparation for Auramine O and Direct Red 23*

Materials

Auramine O (Cas no. 2465-27-2, Sigma Aldrich, <https://www.sigmaaldrich.com/CA/en>)

Direct Red 23 (Cas no. 3441-14-3, Sigma Aldrich, <https://www.sigmaaldrich.com/CA/en>)

Basic Fuchsin (Cas no.569-61-9, Sigma Aldrich, <https://www.sigmaaldrich.com/CA/en>)

Fluorol Yellow 088 (Cas no.81-37-8, ChemCruz, <https://www.scbt.com/p/fluorol-yellow-088-81-37-8>

DMSO

100% Ethanol

Prepared ClearSee

Procedure

1. Make stocks of Auramine O, Direct Red 23, and Basic Fuchsin respectively with ClearSee by adding the correct amount of powdered pigment
   1. 5% (w/v) for Auramine O, 1% (w/v) for Direct Red 23, and 2% (w/v) for Basic Fuchsin
2. Make Fluorol Yellow 088 stock solution by first making a 1% solution of powdered Fluorol Yellow 088 in DMSO,
   1. Use this stock to make a 0.1% solution of dissolved Fluorol Yellow 088 in 100% ethanol, this will be the stock solution that is incorporated into the master mix
3. Wrap tubes in aluminum foil and keep at 4℃, solutions can be kept for several months if properly stored

*Stain master mix preparation*

Materials

5% (w/v) Auramine O

1% (w/v) Direct Red 23

2% (w/v) Basic Fuchsin

5 mM Calcofluor White (Cas no. 4404-43-7, bioWORLD, <https://www.bio-world.com/?zenid=darh23jdjil4nrl6rgtmtviif0>)

Prepared ClearSee

Procedure

1. Option 1 🡪 Working volumes of 0.25mM Calcofluor white, 0.5% (w/v) Auramine O, and 0.1% (w/v) Direct Red
   1. In a 1.5 mL microcentrifuge tube, wrapped with aluminum foil, add 750uL of ClearSee
   2. To the ClearSee, add 100 uL each of the prepared Auramine O and Direct Red 23 stocks and 50 uL of 5mM Calcofluor White, mix stains together by tapping briefly on a vortex
2. Option 2 🡪 Working volumes of 0.0625 mM Calcofluor White, 0.01% (w/v) Fluorol Yellow 088, and 0.2% (w/v) Basic Fuchsin
   1. In a 1.5 mL microcentrifuge tube, wrapped with aluminum foil, add 787.5 uL of ClearSee
   2. To the ClearSee, add 100 uL each of the prepared Fluorol Yellow 088 (ethanol stock) and Basic Fuchsin and 12.5 uL of 5mM Calcofluor White, mix stains together by tapping briefly on a vortex
3. Store stain at room temperature until needed

IMPORTANT TIPS

- Make fresh stain before every intended use
- Modify the amount of master mix prepared dependent on the number of slides you intend to make at a given time
  - 1 tube full of stain master mix is usually enough to stain 4-5 prepared slides

**Block preparation**

Materials

Agarose (Genesee Scientific, Cas no. 90120-36-6, [https://www.geneseesci.com/](https://www.geneseesci.com/?srsltid=AfmBOor8RnoLZhtmbJYmDddSfxlOgdxEXHattUuBP9AVgLafbZxNYo_d))

Sterilized water

Desired tissues/sample

Procedure

1. Make 5% agarose by dissolving the correct amount of agarose in water and microwave until combined
2. Let the agarose cool for a few minutes before adding in a thin layer to your mold
3. Immediately add your desired tissues using forceps to the agarose before it solidifies, taking special care to ensure correct alignment of samples in all directions
4. After the first layer has solidified, gently lift it up from the mold with a stick or the back end of your forceps and place to the side
5. Heat up the agarose again in the microwave
6. Finish by adding a secondary thin layer to the mold before placing the previously solidified layer with the samples on top, yielding a final solidified block with thin layers on each side of the samples

IMPORTANT TIPS

- Be especially careful when microwaving as high concentrations of agarose can be very thick and boil over easily
- We recommend using a square petri dish as a mold because it allows for many samples to be placed at a time and is convenient
- Have your desired samples already cut/prepared and ready for placement, this helps to work quickly and efficiently

**In-slide preparation of sectioned sample**

Materials

A small box

Glass slide with sectioned sample

Kimwipes

Vacuum grease, in a syringe

Prepared fixative

Prepared stain master mix

1x PBS (Thermo Fisher Scientific, <https://www.thermofisher.com/ca/en/home.html>

50% glycerol (Cas no. 56-81-5, Fisher Scientific, <https://www.fishersci.ca/ca/en/home.html>)

Glass coverslips

Procedure

1. Start with a sectioned sample on a glass slide at room temperature
2. Without applying a coverslip, verify the sample using a brightfield microscope, if all structures are intact and the section looks good, proceed with staining
3. Dry off excess water with a kimwipe and label the slide
4. Add vacuum grease using a syringe carefully around the sample, creating a continuous makeshift well, taking care to not add too much
5. Completely cover the sample or fill the well (about 200-400 uL of liquid) with fixative and let it sit for 30 minutes
6. After 30 minutes, use a pipette to gently draw the fixative back up and disperse into a separate tube to be used later
7. Completely cover the sample or fill the well with stain master mix and let the sample sit in a closed box (or in a dark area) for 3 hours, with periodical checks to replenish dried stain
8. During the staining process, agitate or tilt the slide very gently every 15-30 minutes to ensure thorough staining
9. After 3 hours, use a pipette to gently draw the stain back up and disperse into a waste container
10. Wash the sample with 1x PBS 2-3 times with gentle agitation/tilting of the slide to remove excess stain
11. Add 50% glycerol to the well and seal the slide with a coverslip, gently pushing down on the vacuum grease to ensure good contact between the glycerol and the coverslip
12. Store slides in the dark at 4℃ and can be used up to 2-3 weeks after preparation

IMPORTANT TIPS

- Make sure the slide is completely dry before adding vacuum grease, otherwise the grease will not adhere properly to the slide and cause leaking of fixative and stains which is messy/hard to manage
- For all staining and fixing steps, we recommend covering the slide with the lid of a petri dish to avoid spills/accidental touching
- Stains are photosensitive thus we recommend keeping slides in the dark during and after the staining steps
- If samples have moved around during the staining process, you can use a paint brush to reorder them on the slide prior to sealing the slide with 50% glycerol and the coverslip
