## Supplemental table for "In-slide sample preparation allows for rapid triple staining and the detection of specific cell wall components in monocot roots"

|  |  | Total Preparation Time | | |  |  |
| --- | --- | --- | --- | --- | --- | --- |
| Organism | Stains Used | Fixing | Clearing | Staining | Structures and components visualized | Reference |
| White Spruce | Fluorol Yellow 088  Basic Fuchsin Calcofluor White | 1 day | 1-14 days (dependent on organism type) | Dyes dissolved in Clearsee; about 2 hours and 30 minutes (for triple stained samples) | Lignin and suberin in the root endodermis and cellulose in cell walls | Sexauer et al 2021 |
| *Anacyclus clavatus* | Calcofluor White  Basic Fuchsin Auramine O | 1 to 2 hours | For roots, 1 day | 30 minutes to 1 day (dependent on the stain) | General cell wall components including cellulose, lignin, suberin, and xyloglucan | Picinini et al 2024 (original protocol from Urasche et al 2017) |
| Arabidopsis | Calcofluor White Direct Yellow 96  Direct Red 23 Basic Fuchsin Auramine O  Nile Red  Fluorol Yellow 088 | 1 to 2 hours | For roots, 1 day | 30 minutes to 1 day (dependent on the stain) | General cell wall components including cellulose, lignin, suberin, and xyloglucan | Urasche et al 2017 |
|  |  | 1 day | 1-14 days (dependent on organism type) | Dyes dissolved in Clearsee; about 2  hours and 30 minutes (for triple stained samples) | Lignin and suberin in the root endodermis and cellulose in cell walls | Sexauer et al 2021 |
| Broad Bean | Calcofluor White  Basic Fuchsin Auramine O  Fluorol Yellow 088  Safranin Red Astra Blue | 1 to 2 hours | For roots, 1 day | 30 minutes to 1 day (dependent on the stain) | General cell wall components including cellulose, lignin, suberin, and xyloglucan | Picinini et al 2024 (original protocol from Urasche et al 2017) |
|  |  | Samples stored in fixative (no explicit timing mentioned) | N/A | 10 minutes to 1 hour | Lignin and suberin in the root endodermis | Carjaval et al 2025 |
| Cantaloupe | Calcofluor White  Basic Fuchsin Auramine O | 1 to 2 hours | For roots, 1 day | 30 minutes to 1 day (dependent on the stain) | General cell wall components including cellulose, lignin, suberin, and xyloglucan | Picinini et al 2024 (original protocol from Urasche et al 2017) |
| *Lotus japonicus* | Fluorol Yellow 088 Basic Fuchsin Calcofluor White | 1 day | 1-14 days (dependent on organism type) | Dyes dissolved in Clearsee; about 2  hours and 30 minutes (for triple stained samples) | Lignin and suberin in the root endodermis and cellulose in cell walls | Sexauer et al 2021 |
| Mung bean | Fluorol Yellow 088  Safranin Red  Astra Blue | Samples stored in fixative (no explicit timing mentioned) | N/A | 10 minutes to 1 hour | Lignin and suberin in the root endodermis | Carjaval et al 2025 |
| Tomato | Basic Fuchsin Calcofluor White  Fluorol Yellow 088  Aniline Blue  Safranin Red Astra Blue | N/A | Dyes dissolved in Clearsee; 30 minutes to 1 hour (depending on stains used, simultaneous steps) | | Lignin and suberin in the lignified root exodermis and cellulose in cell walls | Manzano et al 2024 |
|  |  | 1 hour | Minimum of 3 days | 30 minutes to 1 hour | Lignin and suberin in the root endodermis | Jo et al 2025 |
|  |  | 1 day | 1-14 days (dependent on organism type) | Dyes dissolved in Clearsee; about 2  hours and 30 minutes (for triple stained samples) | Lignin and suberin in the root endodermis and cellulose in cell walls | Sexauer et al 2021 |
|  |  | Samples stored in fixative (no explicit timing mentioned) | N/A | 10 minutes to 1 hour | Lignin and suberin in the root endodermis | Carjaval et al 2025 |
| Barley | Fluorol Yellow 088  Safranin Red  Astra Blue | Samples stored in fixative (no explicit timing mentioned) | N/A | 10 minutes to 1 hour | Lignin and suberin in the root endodermis | Carjaval et al 2025 |
| Brachypodium | Fluorol Yellow 088 Basic Fuchsin Calcofluor White | 1 day | 1-14 days (dependent on organism type) | Dyes dissolved in Clearsee; about 2  hours and 30 minutes (for triple stained samples) | Lignin and suberin in the root endodermis and cellulose in cell walls | Sexauer et al 2021 |
| Maize | Fluorol Yellow 088  Safranin Red  Astra Blue | Samples stored in fixative (no explicit timing mentioned) | N/A | 10 minutes to 1 hour | Lignin and suberin in the root endodermis | Carjaval et al 2025 |
| Wheat | Fluorol Yellow 088  Safranin Red  Astra Blue | Samples stored in fixative (no explicit timing mentioned) | N/A | 10 minutes to 1 hour | Lignin and suberin in the root endodermis | Carjaval et al 2025 |
